# A Conserved Chromatin-Driven Checkpoint Defines Late Macrophage Maturation Independent of Tissue Specialization

**DOI:** 10.64898/2026.07.01.735834

**Authors:** Djurdja Pasajlic, Maud Plaschka, Frank P. Assen, Anna Kusienicka, Lisa E. Shaw, Peter Traxler, Ulrike Mann, Martin Petrovic, Jelena Bogdanovic, Wolfgang Weninger, Thomas Decker, Florian Halbritter, Matthias Farlik

**Affiliations:** Medical University of Vienna, Department of Dermatology, Vienna, Austria; St. Anna Children’s Cancer Research Institute (CCRI), Vienna, Austria; University of Chicago, Department of Molecular Genetics and Cell Biology, Chicago, Illinois, USA; Max Perutz Labs, Vienna Biocenter Campus (VBC), Vienna, Austria; University of Vienna, Department of Microbiology, Immunobiology and Genetics, Center for Molecular Biology, Vienna, Austria

**Keywords:** Macrophage maturation, innate immune competence, chromatin remodeling, transcriptional memory

## Abstract

Macrophage identity is widely viewed as a product of ontogeny and tissue-specific imprinting. Here, we identify a conserved, chromatin-driven maturation checkpoint that operates independently of initial lineage commitment and broadly across tissue contexts. Using long-term bone marrow-derived macrophage cultures, we uncover a late-stage transition characterized by coordinated transcriptional and epigenomic remodeling. Integration with *in vivo* developmental, tissue-resident, and monocyte-repopulation datasets demonstrates that this program is conserved across ontogenies, tissues, species, and experimental systems, revealing a previously unrecognized stage of macrophage maturation. Functionally, late maturation preserves core macrophage activities while promoting lysosomal expansion and fundamentally rewiring innate immune responsiveness. Mature macrophages display enhanced stimulus-specific responses to interferons and microbial danger signals, coupled to increased metabolic and inflammatory competence while restricting interferon-induced transcriptional memory. Together, our findings identify late macrophage maturation as a conserved regulatory checkpoint that reprograms the logic of innate immune responsiveness through chromatin remodeling shaping innate immune function.

## Introduction

Macrophages are central regulators of tissue biology, contributing to organ development, homeostasis, host defense, and repair. They are present in virtually all tissues as specialized resident populations – including Kupffer cells in the liver, alveolar macrophages in the lung, microglia in the brain, and macrophages in the spleen and gut – where they acquire tissue-specific phenotypes adapted to local microenvironments.^1,2^

Macrophages arise from distinct developmental origins. Early during embryogenesis, yolk sac-derived erythro-myeloid progenitors (EMPs) generate pre-macrophages that seed tissues and give rise to long-lived tissue-resident macrophages (TRMs).^1,3^ Hematopoietic stem cells (HCSs) also arise during embryogenesis and colonise bone marrow before birth. HSCs produce monocytes that can differentiate into macrophages during inflammation or tissue turnover, postnatally.^2^ Evidence from fate-mapping experiments suggests that EMP-derived TRMs have a high self-renewal capacity and require input from monocytes only in very specific situations and tissues.^4^ Accumulating evidence indicates that monocyte-derived macrophages also persist in adult tissues, where they can adopt TRM characteristics and contribute to local macrophage pools, however, the extend of the monocyte-derived contribution varies across organs and has not been fully defined.^5^ For instance, macrophages residing in the gut lamina propria are continuously replenished by monocytes even under steady-state conditions.^6^

Macrophages vary functionally and phenotypically across tissues as a result of the combined influence of ontogeny and the local tissue environement.^7,8^ Initially, macrophage identity is established by lineage-determining transcription factors and concomitant dynamic chromatin remodeling.^9,10^ The pioneering factor PU.1 establishes macrophage-specific enhancer landscapes and cooperates with factors such as IRF8, C/EBP, and KLF family members to drive lineage commitment.^9,11^ Subsequently, tissue-specific transcription factors including GATA6 (peritoneal) and SALLI1 (microglial) drive functional specialization.^7^ Lineage-foundational enhancer landscapes are established during myeloid differentiation, but stay highly plastic, as large-scale epigenetic reprogramming can occur upon environmental changes, with thousands of regulatory elements being reshaped when differentiated macrophages are exposed to new tissue contexts or deprived of existing tissue contexts.^8,9,12–14^ However, while early differentiation and stimulus-driven chromatin remodeling have been extensively characterized, whether and how epigenomic reconfiguration continues during late stages of macrophage differentiation remains poorly understood.

Studying macrophages *ex vivo* is challenging: while primary macrophages can be isolated directly from tissues or generated *in vitro* from monocytes or bone-marrow progenitors under defined cytokine conditions^15,16^, the complex niche-derived signals (including cytokines, stromal interactions, and local cell-cell communication) instructing TRM identities are difficult to reproduce. Multiple studies have shown that splenic macrophages and microglia rapidly undergo transcriptional and phenotypic alterations upon removal from their native environment, reflecting a loss of niche-imprinted programs and adaptation to culture conditions.^13,17–19^ As a pragmatic compromise, bone marrow-derived macrophages (BMDMs) have become the most widely used *in vitro* model, owing to their accessibility, scalability, and genetic manipulability.^20,21^ Discrepancies between transcriptional and epigenomic states of *in vivo* and *in vitro* macrophages are now broadly recognized.^8,14^ However, whether this discrepancy reflects incomplete differentiation, altered maturation trajectories, or fundamental limitations resulting in missing instructive signals of the *in vitro* system remains unclear.

Here, we performed integrated transcriptomic and epigenomic profiling of long-term BMDM cultures. We define a previously underappreciated, chromatin-driven late maturation checkpoint in macrophage development that is conserved across ontogenies and species. By uncoupling maturation from both early differentiation and tissue-specific imprinting, our long-term culture system captures a developmental state that also occurs *in vivo* and is critical for reshaping macrophage responsiveness. Long-term BMDM cultures provides a tractable environment to dissect the regulatory logic of macrophage maturation, and enables systematic interrogation of chromatin remodeling, transcription factor engagement, and regulatory element dynamics that establish macrophage competence independent of tissue specialization.

## Results

### Long-term bone marrow-derived macrophage culture recapitulates a core macrophage maturation program

Bone marrow-derived macrophages (BMDMs) are widely used for functional assays, despite lacking a direct *in vivo* counterpart. Here, we established a long-term *in vitro* culture model that drives BMDMs through transcriptional changes resembling a core macrophage maturation program that is also observed *in vivo*. This model is based on an adapted differentiation protocol that uses both prolonged supplementation and increased concentration of the macrophage colony-stimulating factor (M-CSF) as compared to conventional approaches.^22^ Briefly, BMDMs were differentiated from murine bone marrow aspirate in the presence of M-CSF at a starting dose of 200 ng/mL, which was further supplemented three times during the first 9 days of culture with 100 ng/mL and then omitted thereafter (**Fig. 1A,B**). By day 9, cells exhibited typical macrophage morphology and expressed canonical murine macrophage markers (CD11b and F4/80) at the protein level (**Figs. 1B, S1A, C**), indicating successful monocyte-to-macrophage differentiation. Extended and intensified M-CSF exposure up to day 9 markedly improved the longterm survival and viability of BMDMs in culture compared with conventional 7-day M-CSF-low protocols (**Fig. S1B**).^22^ This provides a robust starting population for functional studies.

**Fig. 1.**
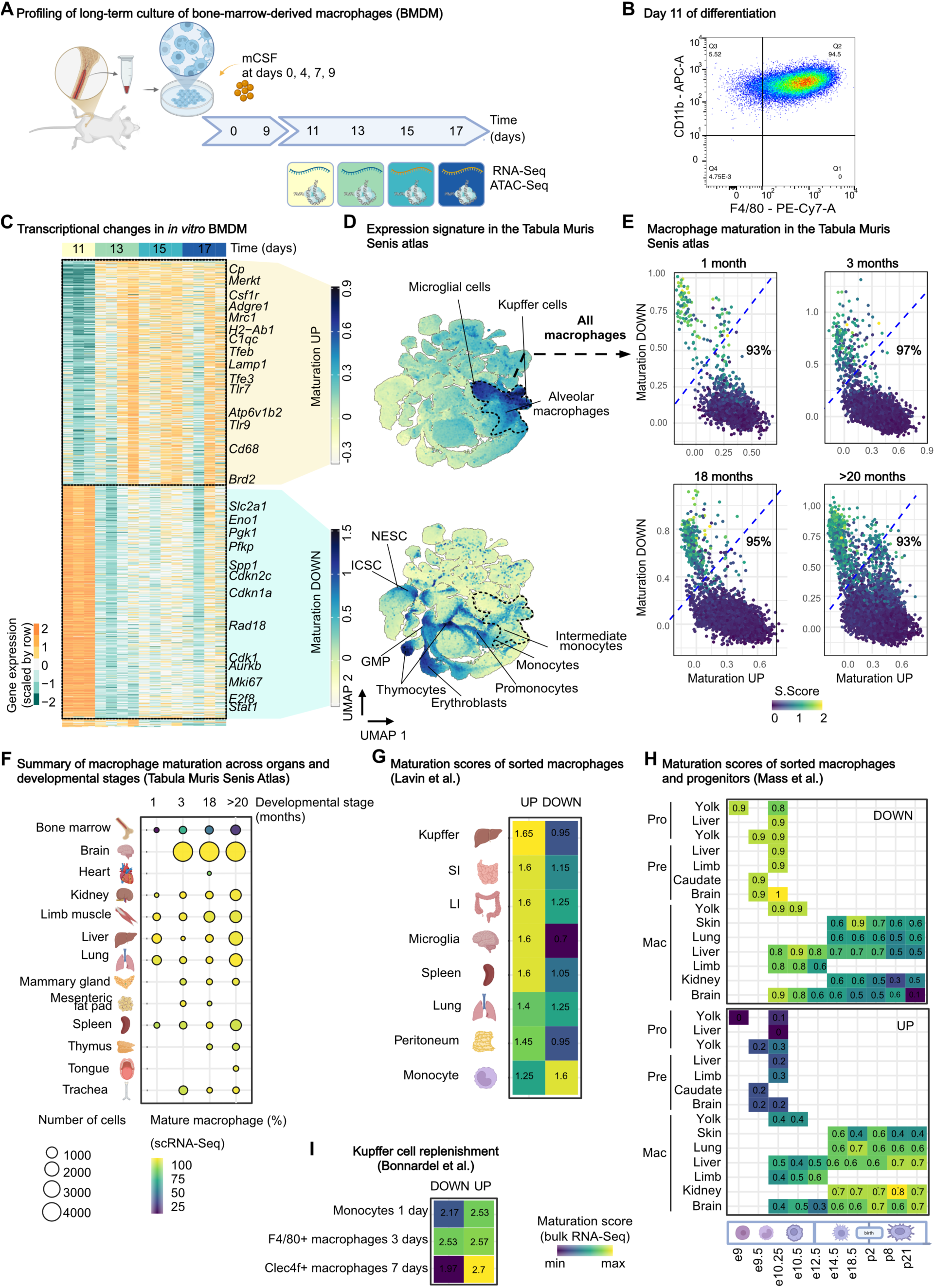
Long-term culture of murine BMDMs recapitulates a core macrophage maturation program with residency-associated features irrespective of origin. A) Schematic of the BMDM differentiation protocol. Bone marrow cells are cultured in DMEM supplemented with M-CSF (200 ng/mL at day 0), refreshed with 100 ng/mL M-CSF at days 4, 7, and 9, and maintained without further media changes until day 17. B) Flow cytometry analysis of canonical murine macrophage markers at day 11 of differentiation. Representative plot shown (gated on live singlets). X-axis: F4/80 (PE-Cy7-A). Y-axis: CD11b (APC-A). 82.8 % of double positive macrophages. C) Heatmap of differentially expressed genes (DEGs) comparing day x + 2 versus day x across late BMDM differentiation (x = 11, 13, 15 and 17 days, n = 3-4 biological replicates per condition). Differential expression was calculated using DESeq2 with apeglm LFC shrinkage (adjusted P < 0.05, |FC| > 1.5). D–F) Re-analysis of the Tabula Muris Senis single-cell RNA-seq dataset. D) UMAP of all cell types, colored by macrophage maturation UP (top) and DOWN (bottom) scores. Scores were computed using AddModuleScore based on DEGs identified between day-13 and day-11 BMDMs. Macrophage clusters are indicated by dashed outlines. NESC, neuroepithelial stem cells; ICSC, intestinal crypt stem cells; GMP, granulocyte-monocyte progenitors. E) Scatter plot of *Maturation-UP* (x-axis) versus *Maturation-DOWN* (y-axis) scores for macrophages at four developmental stages (1, 3, 18, and 20 months). Points are colored by S-phase cell-cycle score. The dashed blue line (x = y + 0.3) empirically separates immature from mature macrophages; percentages of mature cells are indicated. F) Dotplot showing the percentage of mature macrophages across organs (y-axis) and developmental stages (x-axis), defined by high *Maturation-UP* and low *Maturation-DOWN* maturation scores. The size of the dot indicates the number of macrophages in each group and the colors the percentage of mature macrophages. G) Re-analysis of the Lavin et al.^59^ bulk RNA-seq dataset of sorted tissue-resident macrophages and monocytes. *ssGSEA* residency scores were calculated using the BMDM-derived maturation *Maturation-UP* and *Maturation-DOWN* gene sets presented in 1C). H) Re-analysis of the Mass et al.^3^ bulk RNA-seq dataset of sorted pre- and pro-macrophages at different pre- and post-natal developmental stages. *ssGSEA* maturation scores were computed using the *Maturation-UP* and *Maturation-DOWN* signatures established in 1C). I) Re-analysis of the Bonnardel et al.^25^ bulk RNA-seq dataset of bone-marrow-derived macrophages and Kupffer cells following depletion and replenishment. *ssGSEA* maturation scores were calculated using the *Maturation-UP* and *Maturation-DOWN* signatures established in 1C).

RNA-seq revealed that prolonged *in vitro* maintenance induced a pronounced shift in gene expression, occurring abruptly between days 12 and 13 of BMDM culture (**Tables S1, S2)**. Differential expression analysis identified 1,475 differentially expressed genes (DEGs) between 11 and 13 day old macrophages, including 725 upregulated and 750 downregulated transcripts (DESeq2^23^; |log_2_FC| ≥ 0,58, false discovery rate [FDR]-adjusted P-value ≤ 0,05; **Fig. 1C**). Genes downregulated across this transition were enriched for cell cycle, proliferation, and progenitor-associated programs, including markers such as *Mki67*, *Cdkn1a*, and *Cdk1* (**Fig. S1D**). In contrast, upregulated genes reflected a pro-inflammatory, antigen-presenting state, with enrichment of TNFα signaling via NF-κB, complement pathways, and antigen processing and presentation gene sets, alongside increased expression of the macrophage identity marker *Cd68* **(Fig. S1D**). The transcriptional program induced across this transition suggested that these changes indicate a previously unrecognized late-stage maturation process, which occurred robustly across varying culture conditions and could be consistently captured by representative marker genes, with *C1qc* marking the induced state and *Spp1* marking the repressed state (**Fig. S1E**). Notably, the discrete timing of this transcriptional transition indicates that macrophage maturation constitutes a late differentiation event, occurring after establishment of canonical cell-surface marker expression.

To test whether the observed maturation-associated transcriptional program reflects phenomena occurring *in vivo* (rather than being an *in vitro* artifact), we used the previously identified DEGs as gene signatures of maturation and interrogated their expression dynamics in published transcriptomic datasets spanning multiple anatomical locations and embryonic, adult, and aged mice, as well as monocyte-derived and embryo-derived macrophages. We first scored our signatures in the Tabula Muris Senis compendium,^24^ a comprehensive single-cell transcriptomic atlas comprising from 20 mouse organs and tissues. In this unbiased organism-wide dataset, cells with high scores for the downregulated DEGs in our signature (*Maturation-DOWN*) were enriched in progenitors or undifferentiated cell states, including thymocytes, promonocytes, neuroepithelial stem cells, and intestinal crypt stem cells (**Fig. 1D**, lower panel), indicating that this gene set marks early differentiation or progenitor identity. Conversely, cells with high scores for the upregulated DEGs (*Maturation-UP*) were annotated as macrophages (**Fig. 1D**, upper panel). This pattern was consistent across multiple macrophage populations in the atlas, independent of tissue origin, including Kupffer cells, alveolar macrophages, and spleen macrophages. This observation indicates that genes upregulated during long-term *in vitro* culture mirror a core macrophage program conserved across tissues. Next, to determine whether the macrophage-specific enrichment of our signature was influenced by age, we reanalyzed the same dataset focusing on macrophages from different age groups (1, 3, 18, or ≥20 months). Scoring macrophages for the upregulated and downregulated components of our signature revealed that the observed pattern was independent of mouse age: across all four age categories, ≥93% of macrophages exhibited high scores for the upregulated genes and low scores for the downregulated genes (**Fig. 1E**). We next quantified the percentage of mature macrophages within each tissue and age category, revealing uniformly high values across all annotated tissue macrophages at all tested ages (**Fig. 1F**), with the exception of the bone marrow, which contained only a low fraction of macrophages with high maturation scores (25-75%). Thus our data further supports the notion that our signature captures a core and age-independent macrophage maturation program.

To further contextualize our observations in context of known macrophage biology, we utilized three independent datasets. First, we examined data from FACS-purified TRMs from multiple murine tissues.^14^ This analysis confirmed that macrophages across all tissues displayed high scores for our *Maturation-UP* signature, irrespective of tissue origin or specialized phenotype (**Fig. 1G**). Second, we examined data from TRM ontogeny during embryonic development.^3^ Erythro-myeloid progenitors (*pro*) and yolk sac-derived premacrophages (*pre*) exhibited high scores for the downregulated component of our signature (*Maturation-DOWN*) and low scores for the upregulated component (*Maturation-UP*; **Fig. 1H**). As differentiation progressed and macrophages seeded embryonic tissues, this pattern inverted, with mature tissue macrophages displaying high *Maturation-UP* scores and correspondingly low *Maturation-DOWN* scores. Third, we examined macrophage niche repopulation dynamics using a published dataset tracking monocyte-derived replenishment of Kupffer cells (KCs) following depletion (**Figs. 1I, S1F**).^25^ Monocyte-derived KCs exhibited a transient enrichment of the *Maturation-DOWN* signature at day 3 post-depletion, consistent with the reported proliferative expansion and induction of cell cycle-associated programs at this stage (**Fig. 1I**). This signal diminished by day 7. In parallel, the upregulated component of our signature (*Maturation-UP*) was already elevated as early as 24 h after niche entry, coinciding with the rapid loss of monocyte-associated genes and induction of core macrophage and KC-specific transcriptional programs. This score remained stable throughout the proliferative phase and further increased upon acquisition of canonical KC markers such as *Clec4f*.

Together, our analyses solidify the conclusion that the transcriptional switch observed during prolonged BMDM culture reflects a core macrophage maturation program that operates *in vivo* across tissues and developmental contexts.

### The late macrophage maturation program is conserved across murine and human systems

To assess conservation of the core macrophage maturation program in humans, we evaluate the expression of our gene signatures in human single-cell data. We first obtained a comprehensive dataset of human immune cells from the Human Cell Atlas, which comprises *CD45*^+^ immune cells from 14 organs of healthy donors.^26^ As in mice, the *Maturation-UP* score was confined to the myeloid compartment, whereas the *Maturation-DOWN* score was higher in progenitor and proliferating cells (**Fig. 2A, B**). We next examined myeloid heterogeneity and composition within this reference dataset by quantifying the proportion of mature cells across myeloid cell types and tissues (**Fig. 2C**). Mature cells were defined by high expression of the *Maturation-UP* score and low expression of the *Maturation-DOWN* score (cf. previous analysis in mice, **Fig. 1E, F**). Macrophage subclusters 1 (monocyte-derived inflammatory phenotype in heart, liver, and lung, characterized by high expression of *FCN1*) and 2 (transitional monocyte-derived TREM2⁺ FOLR2⁺ macrophages in the heart preceding full lysosomal maturation [CD14⁺ CD68^low^]) comprised few cells with high maturation scores and were predominantly derived from the CD14⁺ monocyte lineage.^26^ In contrast, long-lived TRM subclusters 3 (Notch/RBPJ-associated macrophages in prostate, heart, and esophagus) and 4 (alveolar PPARG⁺ macrophages) consisted entirely of mature cells.^26–28^ Overall, this suggests that macrophage maturation in humans follows similar transcriptional changes as in mice. Importantly, monocytes, DCs, and pDCs exhibited minimal macrophage maturation signal, confirming the specificity of the gene expression signatures, which capture a macrophage-specific late maturation step.

**Fig. 2.**
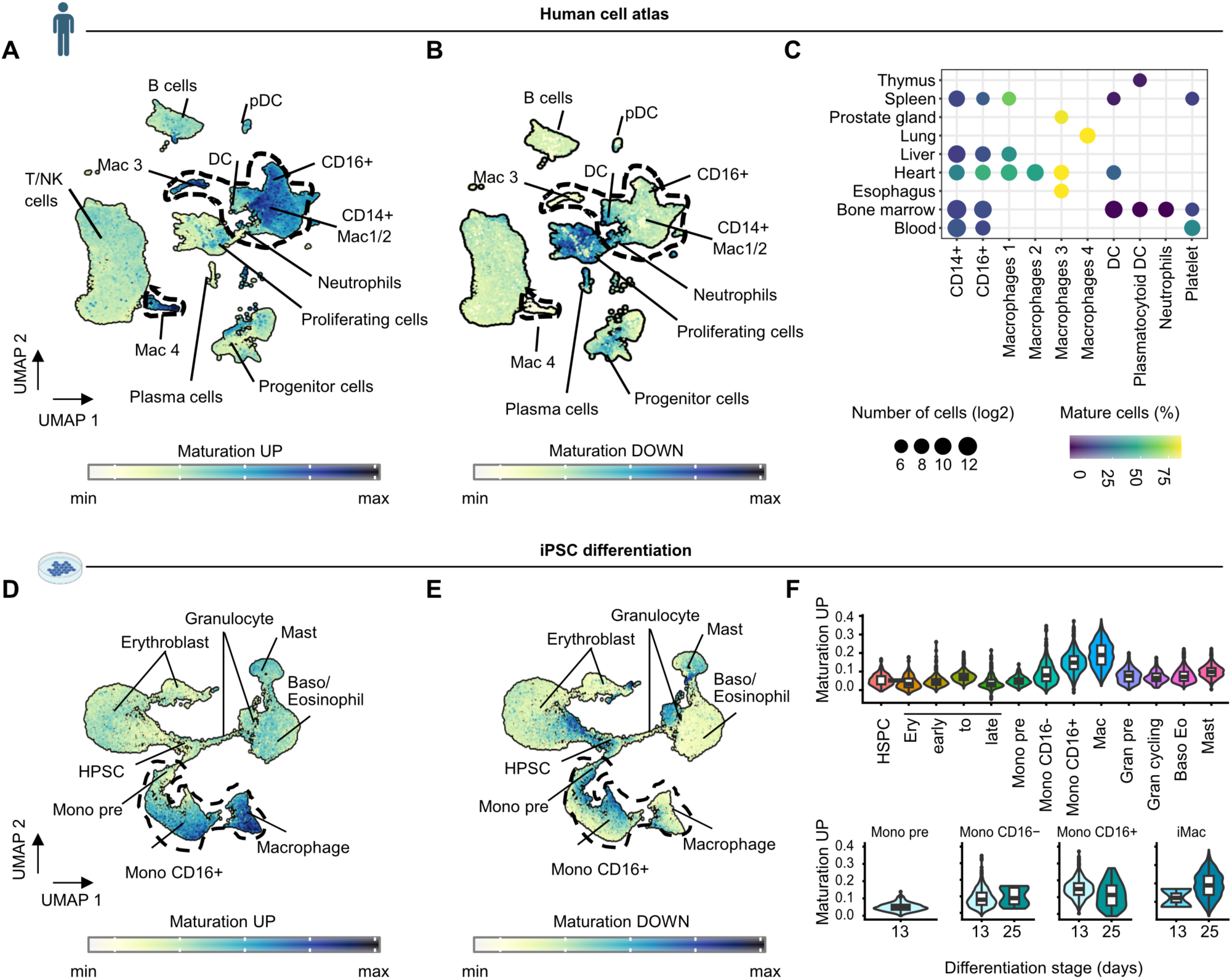
Macrophage maturation is conserved across murine and human systems. A–C) Re-analysis of the human multi-organ CD45⁺ immune cell atlas.^26^ A-B) UMAP of the atlas colored by the macrophage *Maturation-UP* (A) and *Maturation-DOWN* (B) scores. Scores were computed using *AddModuleScore* based on the BMDM-derived maturation gene signatures defined in Figure 1 C). The macrophage cluster is annotated as “Mac”. C) Dot plot showing the percentage of mature myeloid cells across organs (y-axis) and myeloid cell types (x-axis). Macrophage maturation is defined by concomitantly high *Maturation-UP* and low *Maturation-DOWN* maturation scores. Dot size is scaled to the log₂ number of cells per category. D-F) Re-analysis of a human iPSC-derived hematopoiesis single-cell RNA-seq atlas.^29^ D-E) UMAP of the iPSC hematopoietic differentiation model colored by the macrophage *Maturation-UP* (D) and *Maturation-DOWN* (E) scores, calculated using the same BMDM-derived gene sets as in Figure 1C). F) Violin plots of the macrophage *Maturation-UP* score across the full dataset (top; x-axis: annotated cell types) and within the monocyte-macrophage compartment (bottom; x-axis: time after differentiation, in days).

We next asked whether the maturation observed in murine BMDMs was conserved in an human *in vitro* system. To address this, we leveraged data tracking human hematopoietic differentiation from induced pluripotent stem cells (TMOi001-A).^29^ In this dataset, cells differentiating into multiple hematopoietic lineages, including erythrocytes, mast cells, basophils, eosinophils, monocytes, and macrophages, had been collected at two timepoints (day 13 and 25) for scRNA-seq analysis (**Fig. 2D**). We found that the *Maturation-UP* score was highly specific to monocyte-to-macrophage clusters, whereas the *Maturation-DOWN* score was highest in early progenitor populations (**Fig. 2D, E**). The *Maturation-UP* signal increased progressively with time in iPSC-derived macrophages: day 13 macrophages exhibited levels comparable to monocytes, while day 25 macrophages reached the highest maturation scores (**Fig. 2F**).

These findings indicate that macrophage maturation is a time-dependent process conserved across species and independent of progenitor origin, representing a late differentiation step that is uncoupled from initial lineage commitment. Consistent with our *in vivo* analyses, the maturation signature stratifies myeloid populations along a continuum from recently-recruited, monocyte-derived states to fully differentiated TRMs. Importantly, this program is restricted to the monocyte-macrophage axis and is not active in dendritic cell differentiation, suggesting a lineage-specific maturation trajectory.

### Epigenomic remodeling accompanies late macrophage maturation

Having identified a previously unrecognized, macrophage-intrinsic late maturation step that emerges in a temporally defined manner and also occurs *in vivo* across tissues, we next asked whether this transition is accompanied by epigenomic remodeling, which is required for initial lineage specification.^9,30^ As a proxy for epigenome remodeling, we profiled chromatin accessibility using the assay for transposase-accessible chromatin followed by sequencing (ATAC-seq).^31^ We identified 2,162 differentially accessible regions (DARs; DESeq2^23^, |log_2_FC| > log_2_(1.5), FDR-adjusted P-value < 0.05) between immature (day 11) and mature (day 13) macrophages, with the majority (64%) of DARs occurring distal (>3 kb) from the transcription start site of genes (**Fig. 3A, Tables S1, S3**). While the majority of DARs (61%) lost accessibility during maturation, DARs that gained accessibility overlapped significantly with regions previously associated with active histone marks in multiple macrophage populations (e.g., microglia or peritoneal macrophages;^8^ Fisher’s exact test via LOLA^32^, FDR-adjusted P-value < 0.05; **Fig. 3B, Table S4**). These findings indicate that the maturation we observe *in vitro* is not to be considered a passive stabilization phase, but rather a structured and directional process involving both transcriptional changes and active remodeling toward mature macrophage chromatin.^8,12,33,34^ Importantly, BMDMs are conventionally considered fully differentiated by day 6-7 based on morphological and phenotypic criteria.^22,35^ Our data clearly demonstrate that instead a substantial wave of epigenomic maturation occurs considerably later than currently appreciated.

**Fig. 3.**
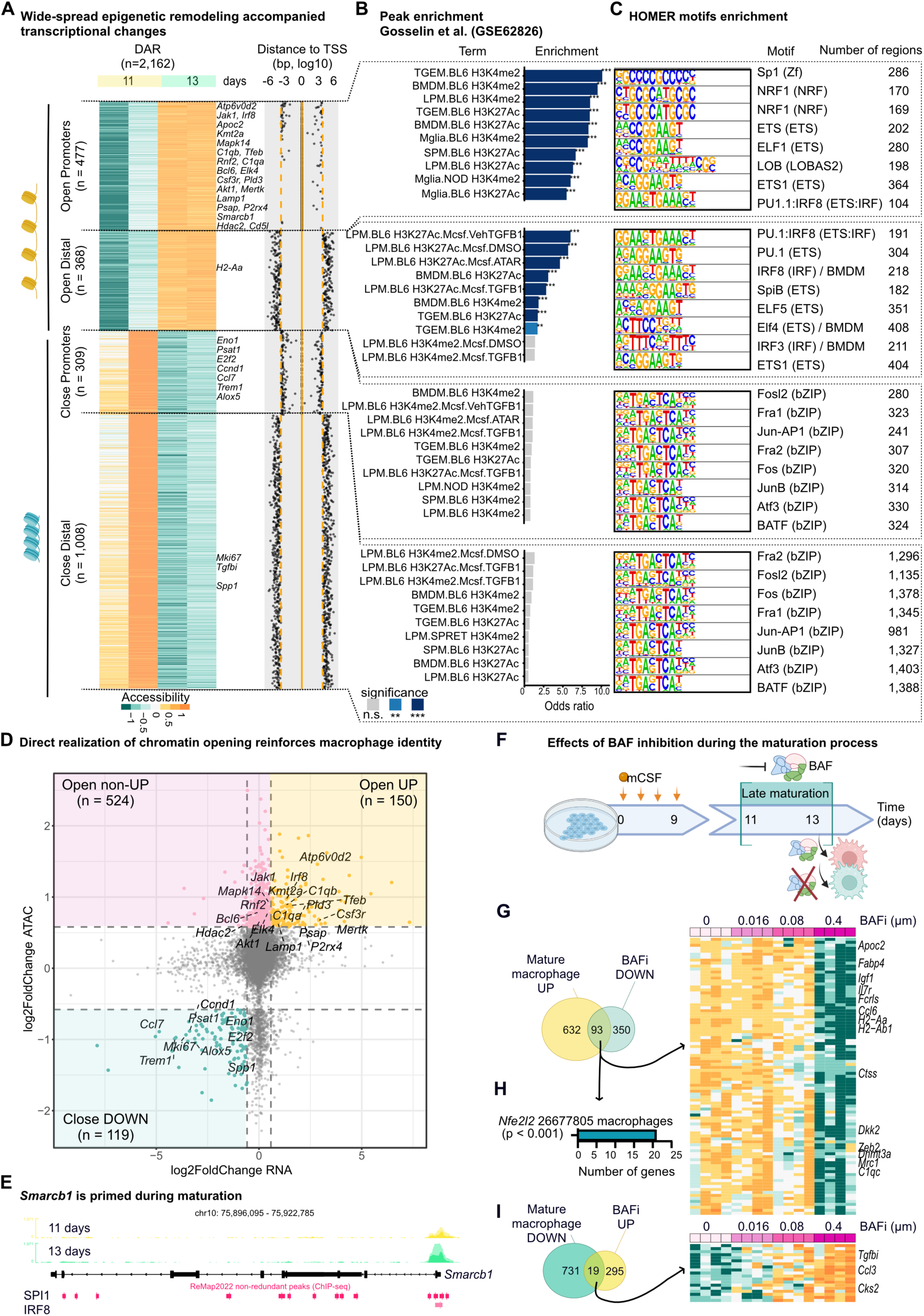
Epigenomic changes during long-term bone marrow–derived macrophage (BMDM) culture. A) Heatmap of differentially accessible regions (DARs) comparing day 13 versus day 11 of the BMDM differentiation protocol (n = 2 replicates per conditions, DESeq2 with *apeglm* LFC shrinkage; *n* = 2,162 DARs; adjusted *P* < 0.05; |fold change| > 1.5). DARs are first ordered according to the log2 fold change (day 13 vs. day 11) and then grouped based on their genomic localization either within promoter regions (±3 kb from the transcription start site, TSS) or outside promoter regions. This allows the definition of four clusters: open promoter (n = 477 DARs), open distal (n = 368 DARs), close promoter (n = 309 DARs) and close distal (n = 1,008 DARs). On the left: Scatter plot indicating, for each DAR shown in the heatmap in A) (y-axis), the distance to the nearest transcription starting site (TSS, x-axis; log10 base pairs from TSS). B) Genomic Locus Overlap Enrichment Analysis (LOLA) of differentially accessible regions (DARs) between day 13 and day 11, stratified according to the four clusters defined in A) (open promoter, n = 477 DARs; open distal, n = 368 DARs; closed promoter, n = 309 DARs; closed distal, n = 1,008 DARs). Enrichment was performed against the catalog of transcriptomes and enhancer landscapes from GSE62826.^8^ Statistical significance was assessed using Fisher’s exact test implemented in LOLA (FDR-adjusted p < 0.05). The x-axis represents the odds ratio, and adjusted FDR values are indicated in the labels (ns, grey; FDR < 0.05, light blue; FDR < 0.005, dark blue). The background universe corresponds to all accessible regions detected in the ATAC-Seq dataset. C) HOMER motif enrichment analysis showing the top transcription factor motifs enriched in open or closed DARs when comparing day 13 versus day 11, stratified by promoter and distal regions. Adjusted *P* value < 0.0005 (Benjamini correction). Number of regions containing the top motifs are indicated on the right. D) Scatter plot displaying the relationship between differential gene expression (DEGs, x-axis; log2 fold change) and chromatin accessibility changes of the nearest gene (DARs, y-axis; log2 fold change) in the comparison of day 13 versus day 11. Regions gaining accessibility and associated with mRNA up-regulation are shown in yellow (n = 150 genes), whereas regions losing accessibility and associated with mRNA down-regulation are shown in blue (n = 119 genes). Region gaining accessibility without increased mRNA expression are shown in pink (n = 524 genes). E) IGV genome browser snapshot of the *Smarcb1* transcription starting site (TSS) locus showing merged ATAC-seq signal from two biological replicates at day 11 (top, turkis) and day 13 (middle, yellow), along with ReMap2022 ChIP-seq binding peaks for SPI1 and IRF8 (bottom). F) Schematic model of the BMDM differentiation protocol highlighting BAF inhibition (BAFi) treatment at day 12 during the late maturation phase. Transcriptomic profiles of BMDMs treated or not with BAFi are compared at day 13. G)-I) Integrated analysis of genes involved in macrophage maturation and modulated by BAF inhibition. G) Venn diagram identifying 93 genes upregulated in mature macrophages (*Maturation-UP*) and downregulated upon BAFi treatment (significant enrichment, hypergeometric test with *phyper* p-value = 4.66 × 10^-28^). Heatmap of the 93 BAFcontrolled genes upregulated in mature macrophages. Gene expression values are rowscaled and displayed from blue (low) to yellow (high). H) Functional enrichment analysis of the 93 genes up-regulated in mature macrophages (*Maturation-UP*) and controlled by BAF using ChEA gene sets.^96,105^ Enrichment was assessed using a one-sided hypergeometric test with false discovery rate correction (Benjamini–Hochberg).^97^ The x-axis shows the number of enriched genes, and adjusted *P* values are indicated in the labels. The background gene universe corresponds to all genes detected in the BAFi RNA-seq dataset. I) Venn diagram identifying genes down-regulated during macrophage maturation (*Maturation-DOWN*) and rescued by BAFi. Heatmap of the 19 BAF-controlled genes down-regulated during maturation. Gene expression values are row-scaled and displayed from blue (low) to yellow (high).

To identify potential regulators of the observed chromatin changes, we next searched the DNA sequences underlying DARs and found motif matches to distinct sets of DNA-binding proteins for each class of DAR (HOMER motif analysis;^33^ **Fig. 3C, Table S5**). We found an enrichment of ETS-family (ETS1, ELF1, SpiB, PU.1) and IRF (IRF8) motifs in regions that became accessible after 13 days of culture (both distal and proximal). Empirical evidence of transcription factor binding in macrophages and related cell types (ChIP-seq data from the ReMap2022 database) corroborated these observations (**Fig. S2A, Tables S4, S6**).^36^ PU.1 is known as a lineage-determining transcription factor in macrophages that pioneers enhancer landscapes and recruits cooperating factors to drive differentiation and functional specialization,^33,37^ while IRF8 modulates PU.1-occupied enhancers to promote macrophage-specific gene programs while repressing alternative myeloid fates.^33,38^ Recent work indicates that PU.1-driven enhancer activation occurs in multiple temporal waves *in vivo*, with later waves marking final specifica-tion.^39^ Our data indicate that prolonged culture is required to induce full epigenetic maturity *in vitro* via new waves of coordinated engagement of PU.1 and IRF8. Conversely, we found that regions that lost accessibility between day 11 and day 13 were significantly enriched for AP-1 family motifs (FOSL2, Fra-1, Jun-AP-1; **Fig. 3C**). This is notable given that AP-1 acts as stimulus-responsive enhancer activator, which engage with lineage factors such as PU.1 in macrophages.^12,34,40^ The selective closure of AP-1-enriched regions therefore suggests progressive shutdown of activation-primed chromatin as macrophages transition toward a stable mature identity.

To decipher whether the identified epigenomic changes had a direct impact on the transcriptomic landscape of mature macrophages, we examined genes occurring in the proximity of each DAR and found a significant overrepresentation of known target genes of IRF8, NFE2L2 (nuclear factor erythroid 2-related factor), and SMARCA4 (SWI/SNF-related, matrix-associated, actin-dependent regulator of chromatin, subfamily A, member 4) (**Fig. S2B, Tables S4, S6**). Integration with our matching RNA-seq data delineated the complex effects on the transcriptional output of maturing macrophages (**Fig. 3A,D, Table S3**): for a first class of genes increased chromatin accessibility yielded direct transcriptional changes. This included the *Irf8* gene itself and other regulators of macrophage identity (*MerTK, Csf3r*) as well as genes mediating core homeostatic functions such as efferocytosis,^41^ including the complement system (*C1qa, C1qb*), and lysosomal biogenesis and phagolysosomal function (*Tfeb, Lamp1, Psap, Atp6v0d2, P2rx4, Pld3*), indicating coordinated activation of the endolysosomal system.^42–44^ This pattern indicates that the late maturation transition reinforces and refines macrophage functions. A second class of genes had gained accessibility but displayed no corresponding change in RNA expression – this was the largest category (n = 538 DARs, 25% of all DARs; **Fig. 3D**). This class encompassed many signaling mediators (*Mapk14*, *Jak1, Akt1*), transcriptional regulators (*Elk4, Bcl6*), and chromatin regulators (*Kmt2a, Hdac2, Rnf2, Smarca1*), suggesting that maturation established a broadly permissive regulatory landscape extending beyond execution of immediate effector programs. While enhancer priming has been extensively described in activated macro-phages,^12,34,45^ our data demonstrate that substantial regulatory accessibility is already intrinsically encoded during maturation itself, prior to external stimulation. Finally, a third class included loci with reduced accessibility paired with reduced transcription, including *E2f2*, *Ccnd1, Mki67, Trem1, Spp1, Ccl7, Alox5, Eno1* and *Psat1*, indicating coordinated shutdown of proliferative, inflammatory-amplifier, and anabolic programs characteristic of immature cells (**Fig. 3D**). In particular, reduced accessibility and expression at *Eno1* and *Psat1*, key components of glycolysis and glycolysis-coupled serine biosynthesis, are consistent with repression of a constitutive growth-associated metabolic state.^46^

Together, our findings demonstrate that macrophage maturation *in vitro* is accompanied by stepwise epigenomic remodelling linked to *Irf8* upregulation, a new wave of PU.1 motif accessibility, and closure of AP-1-enriched regulatory regions. Thus, long-term BMDM cultures are a tractable system to study dynamic lineage-determining transcription factor cooperation and chromatin-based mechanisms governing terminal macrophage maturation.

### BAF activity promotes late-stage macrophage maturation

We demonstrated that late-stage macrophage maturation is accompanied by extensive chromatin remodeling at regulatory elements enriched for lineage-determining transcription factors. Notably, known target genes of the BAF complex component SMARCA4 and BAF binding partner BRD4 were found enriched in opening proximal and distal DARs (**Fig. S2A, B**). The promoter regions of the BAF complex component *Smarcb1* is bound by both macrophage lineage determining factors IRF8 and SPI1 and displayed increased accessibility in day 13 compared to day 11 BMDMs (**Fig. 3E**), suggesting that chromatin remodeling capacity itself may be reinforced during maturation. This would be in line with prior work demonstrating that BAF complexes act downstream of PU.1/SPI1 to stabilize enhancer accessibility in myeloid cells.^47^ We thus hypothesized that BAF activity was dynamically engaged as part of the maturation program.

To directly test this hypothesis, we inhibited BAF activity using a selective BAF inhibitor (BRM/BRG1 ATP Inhibitor-1^48^) at day 12 (immediately prior to the major transcriptional and epigenomic transition) and assessed its impact on macrophage maturation (**Fig. 3F**). BAF inhibition resulted in dose-dependent transcriptional changes with 757 DEGs (DESeq2 ^23^ |log_2_FC| > log_2_(1.5), FDR-adjusted P-value < 0.05, **Fig. S2C**, **Table S7**). Notbably, genes that were downregulated by BAF inhibitor treatment, overlapped significantly with *Maturation-UP* genes (hypergeometric test, P-value = 4.66 × 10^-28^; n = 93 genes in overlap; **Fig. 3G**). This overlap includes regulators of macrophage identity and inflammation like *Zeb2* ^49^, antigen presentation like *H2-Aa*, *H2-Ab1*, *Cd74*, and *Ctss*, innate immune signaling (*Tlr7, Sgk1)*, metabolic and lipid-handling programs (*Plin2*, *Fabp4, Mgll*, *Apoc2*), and tissue residency and homeostasis (*Cd5l* [AIM], *Fcrls*). Conversely, 19 genes that were usually downregulated during maturation (*Maturation-DOWN*) were upregulated by BAFi, including TGF-β target *Tgfbi*, the chemokine *Ccl3* and the cyclin-dependent kinase *Cks2* (**Fig. 3F**). Collectively, these changes indicate that BAF activity is required for full-fledged acquisition of the mature macrophage phenotype.

Interrogation of transcription factor target gene sets in the ChEA database ^50^ identified a significant overlap of NFE2L2 target genes with maturation-associated genes affected by BAF inhibition (21 of 93 genes; **Fig. 3H**).^51^ NFE2L2 (sometimes also called Nrf2, but not to be confused with Nuclear respiratory factor 2 ^52^) acts as a master regulator of cellular antioxidant responses, metabolic reprogramming and macrophage polarization.^53^ Importantly, NFE2L2 can regulate PU.1 (encoded by *Spi1*) expression, as shown in alveolar macrophages.^54^ This supports a model in which BAF cooperates with NFE2L2-driven transcriptional programs to establish the mature macrophage state, integrating lineage-defining transcription factor activity with metabolic and functional specialization programs.

### A late maturation checkpoint rewires macrophage interferon responsiveness and transcriptional memory

We discovered a late checkpoint of macrophage maturation linked to widespread transcriptomic and epigenomic changes. We next sought to assess the functional consequences of this maturation step starting with the defining core functions of macrophages *in vivo*: phagocytosis and lysosomal expansion. First, we measured pHrodo-labeled particle uptake via flow cytometry and found that there was no significant change in phagocytotic capacity throughout the maturation process from day 10 to day 14 of cultured BMDMs (paired t-test, P-value = 0.0622; **Fig. 4A**). Second, we asked whether lysosome-associated functions linked to antigen processing were altered during maturation, as indicated by the enrichment for lysosome-associated genes we found by integrated mRNA and chromatin accessibility analysis (**Fig. 3A, D**). Using β-galactosidase (β-gal) as a surrogate measure for lysosome expansion, we found a significant increase of β-gal-positive cells from <10% at day 10 to >85% by day 14 (paired t-test, P-value = 0.0019; **Fig. 4B**). Importantly, immunofluorescence staining for p16 and p21 confirmed that this increase was not associated with senescence (**Fig. S3A**). Together these findings suggest that while maturation does not affect core phagocytic function, it is accompanied by substantial expansion of the lysosomal compartment, that may be critical to reach full antigen processing capacity.

**Fig. 4.**
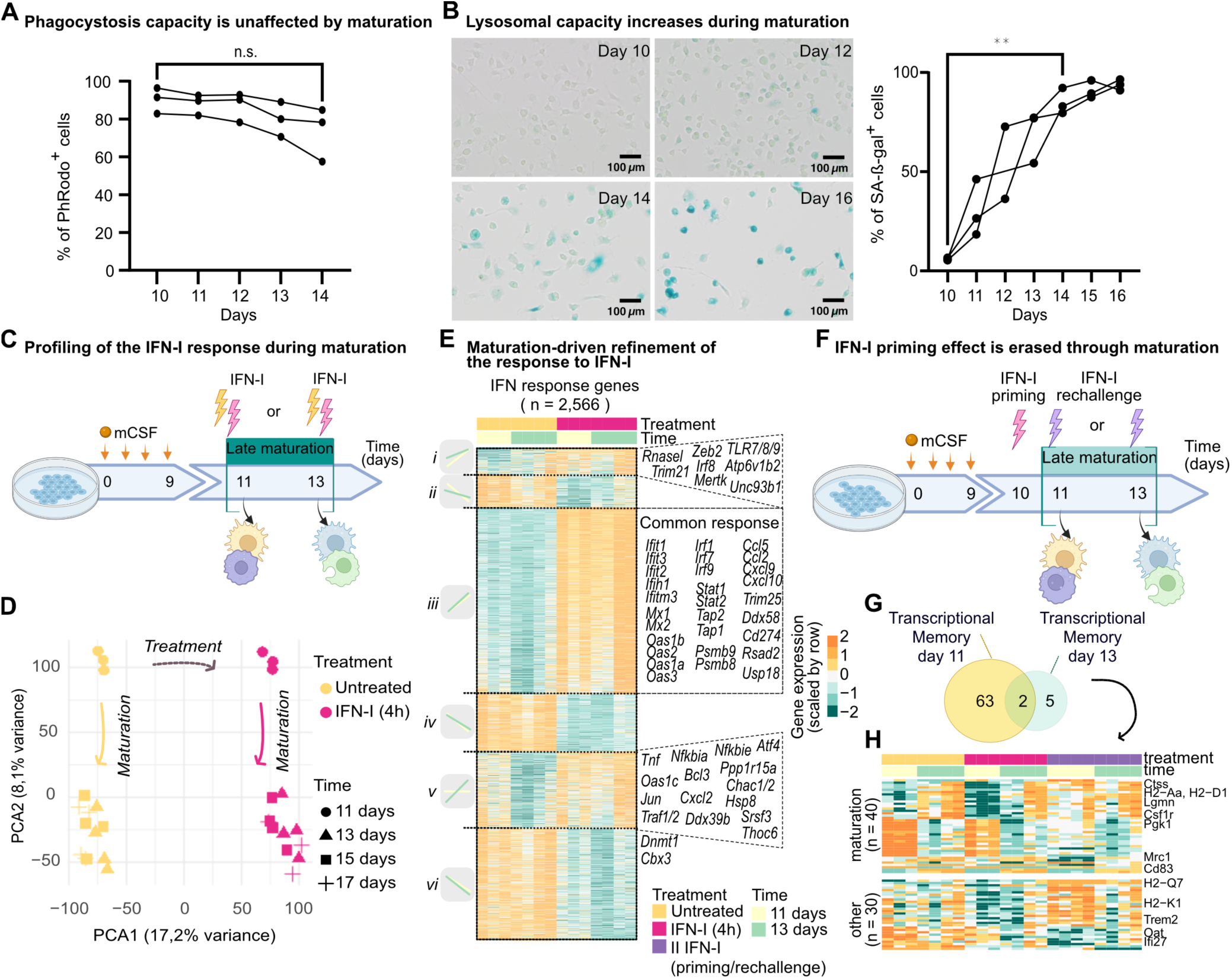
A late maturation checkpoint rewires macrophage interferon responsiveness and transcriptional memory. A) Phagocytosis assay showing the percentage of pHrodo-positive BMDMs following exposure to E. coli fragments over the differentiation protocol (day 10 to 14), measured by flow cytometry. Lines connect paired samples from the same biological replicate (n = 3). Statistical comparison between day 10 and day 14 was performed using a paired two-tailed t test (ns, not significant). B) Senescence-associated β-galactosidase (SA-β-gal) assay showing the percentage of SA-β-gal positive BMDMs from day 10 to day 16, with daily sampling, measured by imaging. Curves represent paired biological replicates (n = 3), each shown as the mean of three technical replicates. Statistical comparison between day 10 and day 14 was performed using a paired two-tailed t test (p-value = 0.0019). Scale bar = 100 µm. C) Schematic of the BMDM differentiation protocol with 4-hour type I interferon (IFN-I) treatment applied at day 11 or day 13. Transcriptomic responses of untreated and IFN-I treated BMDMs were compared between immature (day 11, in yellow) and mature (day 13, in pink) states. D) Principal component analysis (PCA) of untreated (yellow) and IFN-I treated (pink) BMDMs following 4-hour IFN-I stimulation at day 11 (circle), 13 (triangle), 15 (square), or 17 (cross). n = 3-4 biological replicates. Principal component 1 (PCA1, x-axis) reflects treatment effects with 17.2% of variance, whereas principal component 2 (PCA2, y-axis) reflects maturation status with 8.1% of variance. E) Heatmap of differentially expressed genes (DEGs) in response to 4-hour IFN-I treatment in immature (day 11) and mature (day 13) BMDMs (DESeq2 with apeglm LFC shrinkage; adjusted P < 0.05, |fold change| > 1.5). Expression values are scaled by row (from min to max, turquoise to yellow). DEGs are ordered by fold change across conditions. Clusters i-ii represent IFN-I responsive genes specific to immature BMDMs, respectively up and down regulated; clusters iii-iv represent shared IFN-I responsive genes, respectively up and down regulated; clusters v-vi contain IFN-I response genes specific to mature BMDMs. F) Schematic of IFN-I priming and rechallenge experiment. BMDMs were primed with IFN-I at day 10 and rechallenged at day 11 (immature, yellow) or day 13 (mature, green). G) Venn diagram showing overlap of IFN-I memory genes in immature (day 11) and mature (day 13) BMDMs. Memory genes were defined as DEGs uniquely induced upon IFN-I rechallenge relative to a single 4-hour IFN-I treatment and excluding short-term IFN-I response genes. H) Heatmap of expression of the 70 IFN-I memory genes in untreated (yellow), single IFN-I treated (4 h, pink), and rechallenged (purple) BMDMs at day 11 (yellow) and day 13 (green). Expression values are scaled by row (from min to max, turquoise to yellow).

We previously demonstrated that tonic type-I IFN signalling critically contributes to the maintenance of splenic macrophage identity *in vivo.*^13^ To determine how maturation influences cytokine responsiveness, we stimulated macrophages with IFN-β either before (day 11) or after (days 13) establishment of the maturation program for 4 h (**Fig. 4C**). Principal component analysis revealed that IFN-β treatment represented the largest source of transcriptional variation (17.2% variance explained in principal component analysis), whereas the second principal component separated pre- and post-maturation macrophages (8.1% variance; **Fig. 4D**), indicating that maturation substantially alters the cellular response to interferon. Differential expression analysis identified 2,566 differentially expressed genes (**Table S8**), which segregated into six expression clusters defined by their IFN-β responsiveness before (day11) and after (day 13) maturation: clusters (i–ii) comprised genes whose expression was significantly altered by IFN-β only in immature macrophages, clusters (iii–iv) included genes that were equally responsive in both immature and mature states, and clusters (v–vi) contained genes altered significantly exclusively after maturation (**Fig. 4E**). Specifically, clusters (ii), (iv) and (vi) were characterized by genes that display downregulation in response to interferon treatment with different discrete patterns depending on maturation status. Cluster (iii) was characterized by canonical interferon-stimulated genes (ISGs), including *Mx1/2*, *Oas* family members, *Ifit1/2/3*, *Rsad2*, *Isg15*, *Apobec3* and *Zbp1*.^13,55^ Importantly, two clusters revealed maturation-dependent rewiring of interferon responsiveness: Genes in cluster (i) were IFN-inducible in immature macrophages and displayed a trend to be expressed following maturation, indicating that the maturation program activates part of the interferon response independent of external stimulation. These genes included lysosomal acidification factor *Atp6v1b2*, the endosomal TLR trafficking regulator *Unc93b1*, endosomal nucleic-acid sensing receptors *Tlr7/8/9*, and the macrophage regulators *Mertk, Irf8,* and *Zeb2*. Notably, 33% of these genes were found to be significantly upregulated during macrophage maturation, indicating that IFN stimulation of immature macrophages transiently engages components of a maturation-associated program linked to endolysosomal function and pathogen sensing. In contrast, cluster (v), comprising 442 genes, was transcriptionally repressed during maturation yet became selectively inducible by IFN-β only in mature macrophages. Interestingly, this cluster included genes such as *Nf*κ*bia*, *Tnf*, *Traf1*, *Jun*, *Bcl3* and *Cxcl2*, and genes associated with RNA processing, integrated stress response signaling and protein quality control (**Fig. 4E**). Together, these findings indicate that maturation does not substantially alter the canonical interferon response but instead rewires the downstream transcriptional programs engaged by IFN-β, shifting responsiveness from endolysosomal and immune-sensing pathways toward inflammatory, stress-adaptive, and post-transcriptional regulatory programs. Thus, maturation not only alters basal gene expression but also rewires the regulatory architecture of stimulus-responsive genes, selectively priming inflammatory pathways for activation upon interferon exposure.

To further explore the effect of the late maturation regulatory program on effects of gene priming we built on previous discoveries showing that type-I IFNs can prime bone-marrow-derived macrophages for durable recall responses upon re-exposure to the same stimulus.^56^ To test whether macrophages who passed the maturation checkpoint would still show recall response, we re-challenged macrophages that had previously been treated at day 10 with a second 4-h pulse of IFN-β either on day 11 or on day 13 (**Fig. 4F**). We found that 65 genes displayed transcriptional memory if re-challenged at day 11 (**Fig. 4G**, **4H**, **Table S8**). However, at day 13 (after maturation), only 7 genes kept a memory of the earlier stimulation (*Npy, Tmem176a, Ccnb2, Cd83, Myo1e, Sepp1,* and *Hnrnpr*). 32 of the 65 genes displaying transcriptional memory overlap with our *Maturation-UP* gene signature (**Fig. 4H**). Thus, as with the genes in cluster (i) of our previous analysis (**Fig. 4E**) the macrophage intrinsic maturation program uncouples the transcriptional regulation of these genes from stimulus responsiveness.

Together, these data identify macrophage maturation as a regulatory transition that reshapes interferon responsiveness and restricts stimulus-induced transcriptional memory, transforming a plastic, priming-competent state into a more stable and functionally specialized mature state. Having established that maturation rewires cytokine-driven gene regulation, we next asked whether the same principle extends to pathogen-associated danger signals that engage innate immune sensing pathways.

### A late maturation checkpoint establishes stimulus-specific innate immune competence

Given that maturation rewired interferon-responsive gene programs, we next asked whether responses to pathogen-associated molecular patterns (PAMPs) are similarly shaped by maturation status. To address this, macrophages were stimulated on day 11 (pre-maturation) or day 13 (post-maturation) with lipopolysaccharide (LPS), a cognate ligand of TLR4, or poly(I:C), a synthetic mimic of viral double-stranded RNA recognized by TLR3 (**Fig. 5A**).

**Fig. 5.**
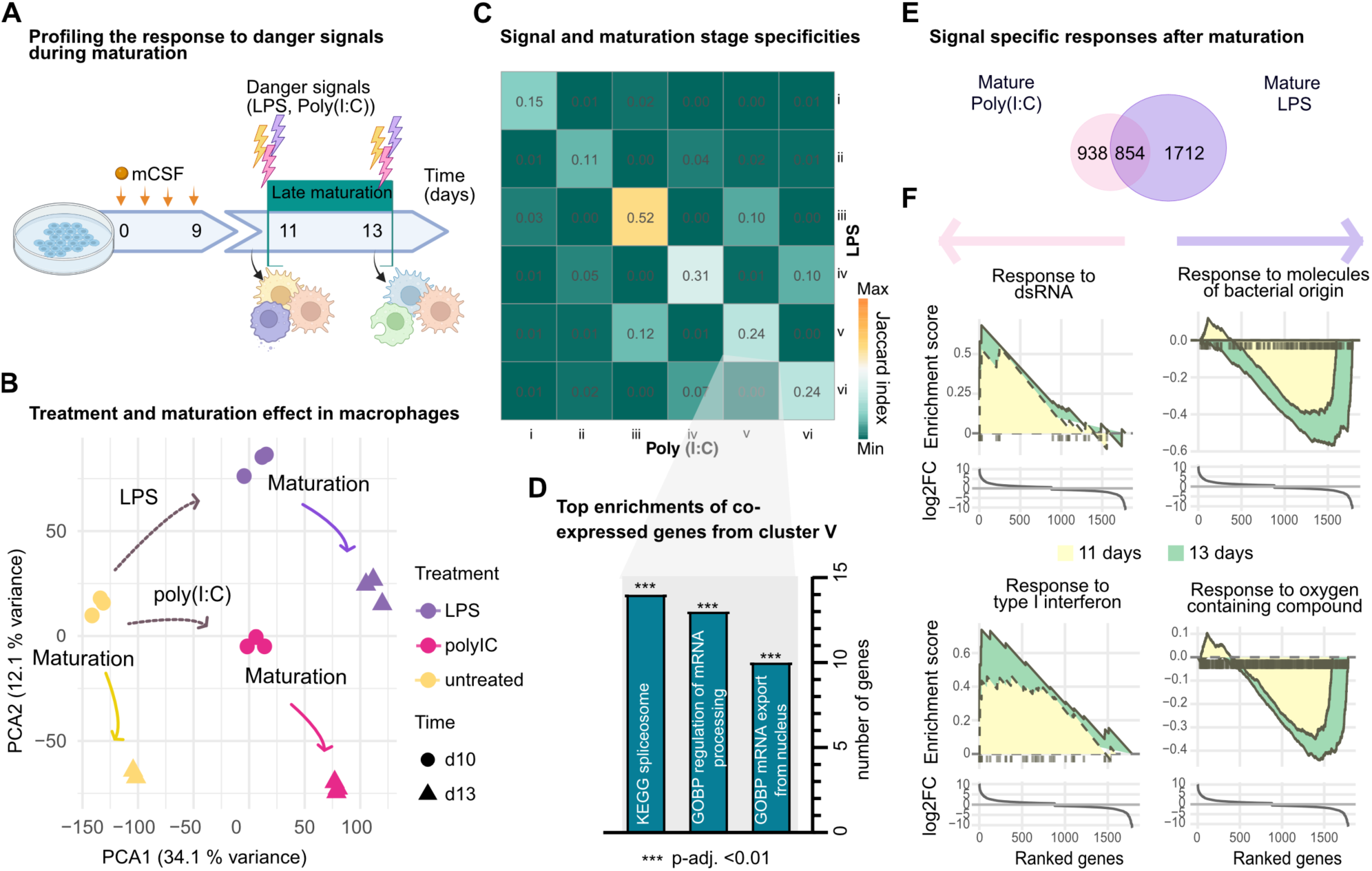
A late maturation checkpoint rewires stimulus-specific innate immune responsiveness. A) Schematic of BMDM stimulation with poly(I:C) (pink) or LPS (purple) at day 11 or day 13, followed by transcriptomic comparison of immature and mature BMDMs. B) Principal component analysis (PCA) of untreated (yellow), poly(I:C)-treated (pink), and LPS-treated (purple) BMDMs at day 11 (circle) or day 13 (square). n = 3 biological replicates, except untreated day 13 (n = 2). Principal component 1 (PCA1, x-axis) captures 34.1% of variance, whereas principal component 2 (PCA2, y-axis) captures 12.1% of variance. C) Heatmap of pairwise Jaccard indices between poly(I:C) and LPS response gene clusters (i–vi, as defined in **Fig. S3**). The Jaccard index was calculated as J(A,B)=∣A∩B∣/∣A∪B∣, where A and B represent gene sets from individual clusters. Continuous scale from minimum to maximum overlap (turquoise to yellow), with higher values indicating greater overlap between gene sets. D) Functional enrichment analysis of the overlapping genes between poly(I:C) and LPS cluster v using ChEA gene sets.^96,105^ Enrichment was assessed using a one-sided hypergeometric test with false discovery rate correction (Benjamini–Hochberg).^97^ The y-axis shows the number of enriched genes, and adjusted *P* values are indicated in the labels. The background gene universe corresponds to all genes detected in the poly(I:C)/LPS RNA-seq dataset. E) Venn diagram showing the overlap between poly(I:C)- and LPS-responsive genes specific to mature BMDMs (i.e. from clusters v and vi). F) Ranked gene set enrichment analysis (GSEA) of selected GO biological processes comparing LPS versus poly(I:C) stimulation at day 11 and day 13. Enrichment curves from day 11 (yellow) and day 13 (green) are shown. For response to double-stranded RNA, enrichment is observed at day 13 (NES = 2.1, FDR = 0.01) but not at day 11. For response to molecules of bacterial origin, enrichment is observed at both day 11 (NES = −1.9, FDR = 0.0075) and day 13 (NES = −2.4, FDR = 4.8 × 10⁻⁶). Similar maturation-dependent differences are observed for type I interferon response and response to oxygen-containing compounds (GOBP response to type I interferon: day 11, no enrichment; day 13, NES = 2.3, FDR = 0.00068. GOBP response to oxygen containing compound: day 11, NES = -1.9, FDR = 1.7 × 10^-5^; day 13, NES = -2.18, FDR = 2.6 × 10^-6^).

PCA revealed that both treatment and maturation status contributed substantially to the observed transcriptional variation (**Fig. 5B**). Poly(I:C) and LPS altered the expression of 4,326 and 5,112 genes, respectively (**Fig. S3B,C, Table S8**). Using the same response-based classification applied to IFN-β stimulation, we defined six maturation-dependent classes of stimulus-responsive genes (**Fig. S3B,C**), indicating that macrophage maturation exerts a consistent influence on transcriptional responses across diverse stimuli. We therefore examined the degree of overlap between corresponding clusters across stimuli. First, a core set of genes responded similarly in both immature and mature macrophages (cluster iii; Jaccard index = 0.52; **Figs. 5C, S3B,C**), indicating that the ability to mount an innate immune response is retained across maturation. These genes included canonical interferon-stimulated and inflammatory regulators such as *Stat1/2, Irf1/7/9, Rel/Nfkb* family members, and antigen presentation machinery (*Psmb8/9/10, Tap1/2, Tapbp*, and MHC class I components), consistent with the induction of endogenous IFN-β downstream of both TLR3 and TLR4 signaling (**Fig. S3D**).

Second, we observed a subset of genes that was preferentially induced in response to both stimuli in immature macrophages only (cluster i; Jaccard index = 0.15; **Figs. 5C, S3B,C**). The same set of genes was already found upregulated by the maturation itself and not further upregulated in response to stimulation. Sixty-one genes were shared between LPS and poly(I:C) responses, including regulators of antigen presentation and immune sensing (*Ciita, H2-Oa, Cd300e, Ms4a* family members), as well as genes linked to membrane trafficking and signaling (*Rab30, Spata13, Akap13*). This pattern suggests that the maturation process by itself mounts a response program biased towards antigen processing capacity and signal modulation, whereas immature macrophages require external stimulation to induce this program.

Third, a maturation-specific inducible program (cluster v) emerged that was shared across both stimuli (Jaccard index = 0.24; **Figs. 5C, S3B,C**). These genes were repressed during maturation but became strongly inducible upon stimulation only in mature macrophages. Notably, this cluster was highly enriched for RNA processing, spliceosome, and nuclear export factors, including *Fmr1, Tpr, Agfg1, Thoc2/6, Rbm8a, Smg7, and Srsf3,* alongside numerous additional components of RNA metabolism and protein homeostasis such *as Sfpq, Sf3b1, Tra2b, Caprin1, Larp1,* and *Psmd6/7* (**Figs. 5D, S3B,C, Table S6**). Thus, maturation appears to actively repress this machinery at steady state while licensing its rapid reactivation upon microbial challenge.

Together, these findings demonstrate that while macrophages retain a shared core inflammatory response irrespective of maturation state, maturation stratifies stimulus-responsive genes into pre-existing (cluster iii), immature-biased (cluster i), and maturation-licensed inducible programs (cluster v). This indicates that maturation does not primarily alter the capacity to initiate inflammatory gene expression, but instead enhances the ability to engage post-transcriptional regulatory machinery, particularly RNA processing and export pathways, thereby enabling more efficient and dynamic transcriptional adaptation upon stimulation.

While these analyses identified maturation-dependent programs shared across microbial stimuli, a substantial fraction of the response remained stimulus-specific. Comparative analysis identified 854 genes shared between LPS and poly(I:C) responses, alongside 938 poly(I:C)-specific genes and 1,712 LPS-specific genes in mature macrophages (**Fig. 5E**, **Table S9**). Ranked gene set enrichment analysis revealed enrichment of antiviral pathways, including response to double-stranded RNA (day 13, NES = 2.1, FDR = 0.01) and type I interferon signaling (NES = 2.3, FDR = 6.8 × 10^−4^) for poly(I:C)-specific genes, whereas LPS-specific genes were enriched for bacterial sensing (NES = −2.4, FDR = 4.8 × 10⁻⁶) and inflammatory metabolism (NES = -2.18, FDR = 2.6 × 10⁻⁶ (**Fig. 5F, Table S10**). Consistent with these enrichments, Poly(I:C) preferentially induced antiviral and nucleic acid sensing programs, including *Ifih1*, *Ddx58*, *Dhx58* and*, Tlr7/8/9*, whereas LPS preferentially activated bacterial sensing and inflammatory pathways, including *Tlr2/6*, *Myd88*, *Ticam1/2*, *Il1a/b*, *Il6*, *Il12a/b* and *Il23a* (**Table S9**). Importantly, these stimulus-specific signatures were detectable at both time points but were stronger following maturation, indicating increased transcriptional specialization of mature macrophages.

Together, these data establish that late-stage macrophage maturation does not primarily alter the capacity to detect danger signals, but instead fundamentally reprograms how macrophages respond to them. Prior to maturation, macrophages exist in a broadly sensing and strongly regulated state that favors signal integration while limiting effector amplification. Following maturation, macrophages acquire highly stimulus-specific transcriptional programs that couple pathogen recognition to inflammatory execution and metabolic adaptation. Combined with the maturation-dependent rewiring of interferon responsiveness and transcriptional memory (**Fig. 4**), these findings reveal that the maturation checkpoint reprograms the logic of innate immune responsiveness, transforming a plastic, priming-competent state into a specialized effector state optimized for tissue immune surveillance and host defense.

## Discussion

Macrophage identity has long been conceptualized as the combined outcome of developmental origin and tissue-specific environmental instruction. Here, we identify a third layer of macrophage biology: a chromatin-driven maturation checkpoint that operates after canonical monocyte-to-macrophage differentiation and independently of tissue specialization. By juxtaposition with transcriptomic data from mouse and human macrophage ontogeny, we demonstrate that this transition represents a broadly conserved maturation state rather than a mere culture-induced phenomenon. Importantly, our findings suggest that macrophage maturation is not simply the terminal phase of differentiation, but a regulatory checkpoint that establishes immune competence by reprogramming how macrophages interpret and respond to environmental signals.

A central implication of our findings is that macrophage maturation can be uncoupled from tissue-specific imprinting. Although local niches are essential to shape tissue-resident macrophage identity,^57^ our data indicate that a substantial component of macrophage competence is established through a cell-intrinsic program shared across tissues and species, thereby extending existing reports of cross-tissues macrophage signatures.^58^ We therefore propose that maturation establishes a competent baseline state onto which tissue-derived cues impose further specialization. This model explains how macrophages can acquire highly tissue-specific phenotypes while retaining a shared core functional architecture.^59^

One exception was the bone marrow, where our analysis suggested macrophages existed in a relatively immature state compared to TRMs. Given the known heterogeneity of the bone marrow comparment,^60^ this may reflect either genuine immature macrophage states or technical limitations of current annotations. Future single-cell profiling of the differentiation trajectory starting from defined bone marrow progenitor populations will be essential to distinguish synchronized maturation from selective expansion of pre-existing subpopulations.

Mechanistically, late-stage macrophage maturation was accompanied by widespread changes in distal chromatin accessibility, consistent with the established role of enhancer remodeling as a major determinant of macrophage cell-state specification.^33^ Accessibility changes were only partly coupled to immediate transcriptional output, suggesting that maturation established regulatory potential required for full functional deployment.^34,45^ The associated motif dynamics support a temporally ordered model: AP-1-enriched regions closed as macrophages exit an activation-primed immature state,^61^ whereas PU.1- and IRF8-associated sites opened during late maturation. This pattern is consistent with established models of iterative PU.1 engagement at regulatory elements and suggests a role for late activity in reinforcing and refining macrophage identity rather than only establishing lineage identity.^62^ Our data further implicate BAF chromatin remodelling as an auxiliary regulator of this checkpoint. BAF inhibition impaired acquisition of maturation-associated programs linked to antigen presentation, metabolic adaptation, tissue residency, and inflammatory competence. These findings extend previous studies implicating BAF chromatin-remodeling complexes in the regulation of macrophage inflammatory responses.^63^ While BAF activity has been shown to control enhancer activation and stimulus-responsive gene expression in activated macrophages, our data suggest that BAF activity also contributes to the establishment of macrophage maturation-associated gene programs, thereby shaping the cellular state from which these responses are executed. Together with the enrichment of PU.1-, IRF8-, and NFE2L2-associated regulatory programs, these data support a model in which BAF acts as a licensing factor that enables the execution of late maturation rather than solely directing initial lineage specification and is consistent with emerging evidence that metabolic state and epigenetic regulation are tightly coupled in mac-rophages.^63–65^ The finding that BAF itself becomes epigenetically engaged during maturation, suggests a feedback mechanism in which chromatin remodeling capacity itself may be reinforced during late differentiation.

Functionally, late-stage maturation does not simply enhance global effector output but instead refines macrophage responsiveness. Phagocytosis remained largely preserved, whereas lysosomal expansion and antigen-processing capacity increased, indicating selective functional tuning. The most profound effect we observed was on stimulus responsiveness: immature macrophages displayed broad sensing capacity, strong negative feedback, and interferon-induced transcriptional memory, consistent with a plastic state optimized for signal integration. Mature macrophages retained canonical IFN and PAMP response pathways but redirected them toward stimulus-specific metabolic adaptation and inflammatory amplification.^46^ In parallel, transcriptional memory was suppressed. In particular, the expression of *Tnf* (encoding TNFα) alongside the inhibitory factors *Nfkbia* and *Nfkbie* argue for an early activation of the TNF-response cascade, which is to our knowledge the first encounter of *Tnf* induction downstream of type-I IFN signaling. Furthermore, an unexpected feature of maturation was the inducibile regulation of the RNA-processing and export machinery in mature macrophages. Genes involved in mRNA maturation, spliceosome function, and nuclear export – including members of the TREX complex (*Thoc2*, *Thoc6*), RNA-binding proteins (*Ythdc*1, *Rbm8a*, *Srsf3*), and nuclear export regulators (*Tpr*, *Agfg1*, *Zc3h11a*) – were transcriptionally repressed at baseline but reactivated following microbial stimulation, suggesting that maturation licensed not only transcriptional responses but also the post-transcriptional infrastructure required to support them. We therefore speculate that maturation optimize the balance between flexibility and execution, ensuring that macrophages respond robustly while maintaining contextual specificity.

Beyond its biological implications, our study has practical consequences for the interpretation of macrophage experiments. Bone marrow-derived macrophages are routinely used as a model of differentiated macrophages and are commonly analyzed within the first week of culture.^22^ Our findings indicate that substantial maturation-associated remodeling continues beyond this period and affects multiple aspects of macrophage behavior, including chromatin organization, stimulus responsiveness, and transcriptional memory. Consequently, macrophage maturation state should be considered an experimental variable alongside genetic background, culture conditions, and activation status when comparing results across studies.

Several questions remain. First, the functional role of chromatin regions that gain accessibility without immediate transcriptional output and their relation to long-term plasticity or context-dependent activation will require testing in future mechanistic studies. Second, while the induction of the maturation program in the *in vitro* culture was robust and reproducible in our hands, the cues that trigger, stablilize, or reverse the onset of the maturation program are currently unknown. Third, single-cell time-course analyses will be required to determine whether maturation reflects a synchronized transition, a selective expansion of subpopulations, or both.

In summary, we identify a conserved, chromatin-driven maturation checkpoint that operates after lineage commitment and independently of tissue specialization. This checkpoint remodels distal chromatin accessibility, engages PU.1/IRF8- and BAF-associated regulatory programs, preserves macrophage core identity, and redefines innate immune responsiveness. By shifting macrophages from a plastic, broadly sensing state towards a stimulus-specific effector state, late maturation acts as a temporal licensing step for immune competence. This conceptual framework and experimental model provides a tractable environment for dissecting how macrophage function is configured across development, homeostasis, and disease.

## Methods

### Isolation of cells from bone marrow

We followed the protocol described by Taxler et al..^66^ Femurs and tibias were obtained from 8-12 week old Cas9 mice (Jackson Laboratory, cat. no. 024858), cleaned of surrounding tissue, and crushed in PBS using a sterile mortar and pestle. Bone marrow containing supernatant was filtered through 100 μm and 45 μm cell strainers. Cells were counted, centrifuged at 4°C for 5 minutes at 400 rcf and resuspended in DMEM-GlutaMAX (Gibco, cat. no. 31966047) with 40% FBS. The cell suspension was mixed with an equal volume of DMEM GlutaMAX with 40% FBS and 30% DMSO (Sigma, cat. no. 1019001000), frozen at 50 million cells/ml, and aliquots were stored at -80 °C (short-term) or in liquid nitrogen (long-term storage).

### Establishment of BMDM

Bone marrow derived macrophages were differentiated by culturing 1 × 10⁸ cells of Cas9 bone marrow in 20 mL of complete media (DMEM (Gibco, cat. no. 41966-029) with 10% FBS and 1% penicillin-streptomycin (Gibco, cat. no. 15140122)) with 200 ng/mL M-CSF (Peprotech, cat. no. 315-02). On day 4 and day 7, 5 mL of the cell culture medium was discarded and 5 mL of fresh DMEM containing 400 ng/mL M-CSF (100 ng/mL final concentration in 20 mL) was added. On day 9, cells were scraped, centrifuged at room temperature for 5 min at 350 rcf, counted and seeded for further experiments. Seeding was done at density of 0.5 × 10⁶ cells/mL, in DMEM with 100 ng/mL of M-CSF per well in a single 35 mm dish or 6 well plate. All cell culture work was done in tissue culture non treated dishes.

### Experimental design

Unless otherwise stated, all experiments were performed using three biological replicates derived from independently harvested bone marrow samples obtained from distinct mouse litters. Cells from each biological replicate were differentiated and processed independently throughout the experiment.

### FACS characterization

Cells were harvested, washed with PBS, pelleted by centrifugation (2,200 rpm, 5 min, 4 °C), and resuspended in autoMACS Running Buffer (Miltenyi Biotec, cat. no. 130-091-221) containing Fc receptor blocking antibody (anti-CD16/CD32; BD Pharmingen, clone 2.4G2, cat. no. 553141). Cells were stained with fluorophore-conjugated antibodies for 30 min at 4 °C, washed, filtered through a 40-µm strainer, and incubated with DAPI for dead-cell exclusion. Samples were analyzed on a FACSAria II flow cytometer (BD Biosciences), and data were processed using FlowJo software (BD). Antibodies used were: anti-CD45 PerCP/Cy5.5 (BioLegend, cat. no. 103132), anti-CD11b APC (BioLegend, cat. no. 101212), anti-F4/80 PE/Cy7 (BioLegend, cat. no. 123114). Debris and doublets were excluded based on forward- and side-scatter characteristics. Macrophages were defined as live CD45⁺ CD11b⁺ F4/80⁺ cells.

### Quantitatve RT-PCR

Total RNA was isolated using the RNeasy Mini Kit (Qiagen, cat. no. 74104) according to the manufacturer’s instructions. Concentration of isolated RNA was measured using a NanoDrop instrument (Thermo Fisher). cDNA was synthesized from 200 ng of total RNA using LunaScript^®^ RT SuperMix Kit (New England Biolabs, cat. no. E3010L). Quantitative PCR was performed using 2 ng of cDNA and GoTaq qPCR Master Mix (Promega cat. No. A6002) according to vendor’s protocol, on a Bio-Rad CFX Opus 96 machine. Primers used were: C1qc_FW CCAAGGGAGAGCCAGGAATC; C1qc_REV GTTTGTATCGGCCCTCCACA; Spp1_FW GGCTGAATTCTGAGGGACTAA; Spp1_REV ATCTGGGTGCAGGCTGTAAA, Gapdh_FW GGGGTCCCAGCTTAGGTTCA; Gapdh_REV CCCAATACGGCCAAATCCGT. The qPCR reactions were run in a technical duplicate with the following settings: 2 minutes activation at 95 °C followed by 40 cycles of 15 seconds at 95 °C, 1 minute at 60 °C (for *Spp1* and *Gapdh*) or 65°C (for *C1qc*) and a melt curve. Cq values were determined using the default settings of the Bio-Rad CFX Maestro software version 2.3 (5.3.022.1030). Relative gene expression was calculated using the ΔΔCt method with *Gapdh* as the reference gene.

### Phagocytosis assay

Phagocytic activity was assessed using pHrodo™ Deep Red–labeled beads (Thermo Fisher Scientific, cat. no. P35361). Beads were resuspended at 2 mg in 2 mL PBS (1 mg/mL). For each condition, 1×10⁶ macrophages in 1 mL of complete medium was treated with 100 µL of the bead suspension and mixed gently. Cells were incubated for 30 min at 37 °C to allow phagocytosis. The reaction was terminated by placing samples on ice, followed by immediate harvesting of the cells. Phagocytic uptake was quantified by flow cytometry based on pHrodo Deep Red fluorescence, which increases upon internalization and acidification within phagosomes. Data was analyzed as the percentage of pHrodo-positive cells.

### SA-ß-galactosidase assay

Senescence-associated β-galactosidase staining was performed using the Senescence β-Galactosidase Staining Kit (Cell Signaling Technology, #9860) following the manufacturer’s protocol. Briefly, cells were washed with PBS and fixed using the supplied fixative solution at room temperature. After fixation, cells were incubated with freshly prepared β-galactosidase staining solution (pH 6.0) at 37 °C in a CO₂-free incubator overnight. β-galactosidase staining was visualized by bright-field microscopy using an Olympus IX83 inverted microscope and 10X objective captured with a color camera (UC90 Olympus). The images were analyzed using a custom Python pipeline, applying white-balanced using a user-defined background region, after which individual cells were segmented with the Cellpose cyto3 model.^67^ For each segmented cell, RGB intensity- and morphology statistics were extracted and classified as senescence-positive (high and low positive, grouped together for downstream analysis) or -negative using a random-forest classifier, that was user-trained on a small subset of data. The trained classifier was then applied across all datasets to quantify over 97,000 cells from 7 culture time points (days 10–16) and three biological replicates per time point.

### Immunofluorescent staining for p21 and p16

Cells (1 × 10⁶) were seeded onto 35-mm untreated polymer coverslip-bottom microscopy dishes (Ibidi, cat. no. 81151) and fixed in 4% paraformaldehyde for 15 min at room temperature. After washing twice with PBS, cells were permeabilized for 30 min in immunostaining buffer (IBS) prepared as previously described in ^68^ and blocked for 30 min in IBS containing 1:100 donkey serum (Sigma-Aldrich, cat. no. D9663). Cells were incubated with primary antibodies against p21 (1:250; Abcam, cat. no. ab107099) and p16^INK4a (1:100; Abcam, cat. no. ab108349) for 1 h at 4 °C, washed three times with PBS, and incubated with Alexa Fluor–conjugated secondary antibodies (donkey anti-rat Alexa Fluor 647 (Jackson ImmunoResearch, code: 712-605-150) and donkey anti-rabbit Alexa Fluor 555 (Jackson ImmunoResearch, code: 711-565-152); 1:400) for 1 h at 4 °C. Nuclei were counterstained with DAPI, and samples were mounted using Ibidi mounting medium (Ibidi, cat. no. 50001). Images were acquired on a Zeiss LSM 900 inverted confocal microscope and 10X objective, using identical acquisition settings for all samples within each biological replicate. For each condition and biological replicate, three images were acquired from distinct regions of the culture dish to capture spatial variability across the culture and provide a representative sampling of cells. Images were then analyzed to classify individual nuclei as negative, P16-positive or P21-positive. Nuclei were segmented from the DAPI channel using the Cellpose cyto3 model^67^, and object classification was performed in ilastik^69^ using the nuclear masks and corresponding raw P16/P21 image data. The resulting object-prediction images, in which each class was encoded by a unique label intensity, were processed with a custom Python script to count and export the number of cells assigned to each class.

### LPS, Poly-IC stimulation

Cells were seeded at 0.5 × 10⁶ cells/mL in complete medium containing 100 ng/mL M-CSF in six-well plates on day 9. The following day, cells were stimulated with LPS at a final concentration of 10 ng/mL (Merck, cat. no. L2630) or poly(I:C) (HMW) at a final concentration of 10 µg/mL (InvivoGen, cat. no. tlrl-pic) for 4 h. Both treatments included matched PBS-treated negative controls. After completion of treatments, cells were washed with PBS, harvested by scraping in 1 mL PBS per well, pelleted by centrifugation (2,200 rpm, 5 min, 4 °C), lysed in 350 µL RLT buffer (Qiagen RNeasy Kit) supplemented with 1% β-mercaptoethanol, and stored at −80 °C until RNA extraction.

### IFN-I priming and restimulation assay

Cells were seeded at 0.5 × 10⁶ cells/mL in complete medium containing 100 ng/mL M-CSF in six-well plates on day 9. The following day, cells were treated with recombinant IFN-β at a final concentration of 1,000 U/mL (PBL Assay Science, cat. no. 12401-1) for 4 h, after which cells were washed once with PBS and replenished with fresh complete medium containing 100 ng/mL M-CSF. After 24 h, cells received a second IFN-β treatment (or a first treatment for control conditions). All treatments included matched PBS-treated negative controls. After completion of treatments, cells were washed with PBS, harvested by scraping in 1 mL PBS per well, pelleted by centrifugation (2,200 rpm, 5 min, 4 °C), lysed in 350 µL RLT buffer (Qiagen RNeasy Kit) supplemented with 1% β-mercaptoethanol, and stored at −80 °C until RNA extraction.

### BAF inhibition

Cells were seeded in six-well plates on day 9 of differentiation and left untreated until day 12. On day 12, cells were treated for 24 h with the BRM/BRG1 ATPase (SWI/SNF) inhibitor BRM014 (MedChemExpress, cat. no. HY-119374) at final concentrations of 400, 80, or 16 nM, with PBS-treated wells included as controls. Following treatment, cells were washed with PBS, harvested by scraping in 1 mL PBS per well, pelleted by centrifugation (2,200 rpm, 5 min, 4 °C), lysed in 350 µL RLT buffer (Qiagen RNeasy Kit) supplemented with 1% β-mercaptoethanol, and stored at −80 °C until RNA extraction.

### RNA-seq data generation

Bulk RNA-seq was performed as previously described using the Smart-seq2 protocol.^13,70^ Total RNA was isolated using the RNeasy Mini Kit (Qiagen, cat. no. 74104) according to the manufacturer’s instructions and eluted in 30 µL RNase-free water. RNA concentration was measured using the Qubit™ RNA HS Assay Kit (Thermo Fisher Scientific, cat. no. Q32854). For cDNA synthesis, 400 pg of total RNA was used. cDNA concentration was determined using the Qubit™ dsDNA HS Assay Kit (Invitrogen), and sample quality was assessed using an Agilent TapeStation with High Sensitivity D5000 ScreenTape (Agilent Technologies, cat. no. 5067-5592). cDNA was diluted to 0.1 ng/µL, and 5 µL was used for library preparation with Nextera XT index adapters (Illumina, Nextera® XT Index Kit v2, Set A). Library quality was confirmed for a subset of samples using D1000 ScreenTape (Agilent Technologies, cat. no. 5067-5582). Per batch, libraries were pooled to 4 nM and sequenced on an Illumina NovaSeq X 100 cylces flow cell in 50 bp paired-end configuration. An overview of all genomic datasets generated in this study is available in **Table S1**.

### Preprocessing of RNA-seq data

After converting BAM files to FASTQ using samtools v1.20,^71^ raw sequencing reads were processed using the nf-core/rnaseq pipeline (version 3.14.0) implemented in Nextflow (version 23.10.1).^72–75^ Unless otherwise specified, all steps were executed with default pipeline parameters. Initial quality control was performed with FastQC (v0.12.1),^76^ and adapter trimming was carried out using TrimGalore (v0.6.7) with Cutadapt (v3.4). Reads were aligned to the iGenomes Mus musculus GRCm38 reference genome with Ensembl gene annotation release 81 using STAR (v2.6.1d) ^77^ for splice-aware mapping. Transcript-level quantification was performed with Salmon (v1.10.1),^78^ and gene-level counts were generated using featureCounts from the Subread package (v2.0.1).^79^ Post-alignment quality metrics were assessed using Qualimap (v2.3) ^80^ and RSeQC (v5.0.2).^81^ Duplicate reads were marked using Picard (v3.0.0), and duplication rates were further evaluated with dupRadar (v1.28.0). Additional QC summaries were compiled with MultiQC.^82^ The pipeline also included transcriptome indexing with RSEM (v1.3.1) ^83^ and STAR (v2.7.10a), ^77^ as well as auxiliary steps for GTF processing and chromosome size retrieval.

### ATAC-seq data generation

ATAC-seq libraries were generated according to the protocol described by Taxler et al.,^66^ without modification. Briefly, 5 × 10⁴ cells were lysed and subjected to Tn5-mediated tagmentation according to the published protocol. Fragmented DNA was purified using the MinElute PCR Purification Kit (Qiagen, cat. no. 28006). Library amplification was performed using indexed ATAC-seq primers,^31^ with the number of enrichment cycles determined by quantitative PCR to avoid overamplification. Libraries were purified by double-sided size selection using AMPure XP beads (Beckman Coulter, cat. no. A63881). DNA concentration was determined using the Qubit dsDNA HS Assay Kit (Thermo Fisher Scientific, cat. no. Q32854), and library molarity was estimated using a High Sensitivity DNA Bioanalyzer chip (Agilent, cat. no. 5067-4626). Libraries were dilute to 4 nM, pooled and sequenced on an Illumina NovaSeq SP platform with 100 cycles in paired-end 50-bp configuration at the Biomedical Sequencing Facility, CeMM. An overview of all genomic datasets generated in this study is available in **Table S1**.

### Preprocessing of ATAC-seq data

After converting BAM files to FASTQ using samtools v1.20,^71^ reads were processed with the nf-core/atacseq v2.0 pipeline (Nextflow v23.10.1) ^72–75^. Unless otherwise specified, all steps were executed with default pipeline parameters. Adapter and quality trimming were performed with Trim Galore v0.6.7 (cutadapt v3.4), followed by quality control with FastQC v0.11.9 ^76^ and MultiQC.^82^ Reads were aligned to the iGenomes Mus_musculus GRCm38 reference (Ensembl gene annotation release 81) using BWA-MEM v0.7.17;^84^ secondary processing included sorting and indexing with samtools (v1.16.1) ^71^ and duplicate marking with Picard v2.27.4.^85^ Blacklist filtering and mitochondrial-read handling followed the nf-core defaults. Peak calling was performed with MACS2 v2.2.7.1, using an effective genome size (--gsize) of 2,652,783,500 for mouse.^86^ Normalized coverage tracks (Big-Wig) and QC metrics were generated with deepTools v3.5.1,^87^ ataqv v1.3.0 (including TSS enrichment), and FRiP/other metrics via bedtools v2.30.0;^88^ summary reports were compiled with MultiQC. The workflow was executed in a containerized environment using Podman, ensuring full reproducibility of software versions and dependencies (see Data and code availability).

### RNA-seq and ATAC-seq data analysis

RNA-seq and ATAC-seq analyses were performed in a containerized environment using Podman to ensure full reproducibility of software versions and dependencies. Briefly, the RStudio Server (R v4.4.1) was built from the based bioconductor_docker:3.19 image.^89,90^ Package versions and dependencies were managed using renv (v1.1.4).^91^ For RNA-seq, the STAR Salmon output ‘salmon.merged.gene_counts.rds” was used. Low expressed genes were filtered in two steps: (i) using Count Per Million (CPM), we removed genes that did not reach a CPM value of 0.3 in at least 3 samples (out of 15 total) and (ii) genes with no more than 10 reads in at least 3 samples. For ATAC-seq, we used the consensus peaks from MACS2 ‘consensus_peaks.mLb.clN.featureCounts.txt”. We removed peaks that did not reach 50 reads in at least one sample (out of 4 samples total) and worked with the remaining 70,322 peaks. Remaining peaks were annotated using *annotatePeak* from ChIPseeker (v.1.42.1).^92^ For both ATAC-seq and RNA-seq, differential expression analysis was performed using DESeq2 (v.1.44.0)^23^ build on the filtered raw counts with a design built on a unique variable combining the time and treatment condition when applicable. Log2 fold changes (LFC) were shrunken using *lfcShrink* type *apeglm* (v.1.28.0).^93^ Genes or peaks with a fold change > 1.5 and an adjusted p-value < 0.05 were considered as differentially expressed genes (DEG) or differentially accessible regions (DAR) respectively. We used the enricher function (default parameters) from the clusterProfiler package (v.4.14.6)^94^ to perform hypergeometric tests for functional enrichment analysis. The universe/background was defined as the row names of the dds object (that is, all detected genes or associated regions). The human hallmark and Gene Ontology (GO) biological process (C5:BP) gene sets were retrieved from the Molecular Signatures Database (MSigDB) using the msigdbr function and package (v.24.1.0)^95^ and the ChEA_2022 gene set was downloaded from https://maayanlab.cloud/Enrichr/#libraries.^96^ Finally, we applied the Benjamini-Hochberg method to control false discoveries in multiple hypothesis testing.^97^ Plots were generated using pheatmap (v.1.0.12),^98^ ggplot2 (v.3.5.2)^99^ ggVennDiagram (v.1.5.2)^100^ and RColorBrewer (v.1.1-3; http://www.ColorBrewer.org) packages.

### Signature mapping to publicly available datasets

Differentially expressed genes (DEGs) in BMDM maintained in culture for 13 days versus 11 days (adjusted p-value < 0.05, |logFoldChange| > 1.5) were used to calculate maturation scores up (logFoldChange > 0) and down (logFoldChange < 0). For bulk RNA-seq data, we used the non-parametric, unsupervised Gene Set Enrichment Analysis (GSVA, v2.0.7, with the functions ssgseaParam and gsva) ^101^ to estimate the variations of the up and down maturation scores though the datasets. Enrichment scores were plotted as heatmap using ggplot2 (v.3.5.2) ^99^ with viridis color maps.^102^ For single-cell data, module scores were calculated using the Seurat function *AddModuleScore* (v5.3.0).^103^ Scatter plots were used to visualize the inverse correlation of the maturation score down (y) and maturation score up (x). In each dataset, a linear model y = x + *intercept* was used to separate mature (y < x + *intercept*) from early (y > x + *intercept*) macrophages and estimate the percentage of mature macrophages per category. For the analysis of the Tabula Muris dataset, *intercept* = 0.3, while *intercept* = -0,1 for the analysis of the Human Cell Atlas dataset. Tables were handled using dplyr (v1.1.4)^104^ Percentages of mature macrophages were plotted as heatmap or dotplot using ggplot2 (v.3.5.2) ^99^ with viridis color maps.^102^

### HOMER motif enrichment analysis

For each gene, the promoter region was defined as ±3,000 bp relative to the transcription start site (TSS). DARs were classified into four categories based on accessibility change (open or closed) and genomic location (promoter or non-promoter). Motif enrichment analysis was performed separately for each DAR category using HOMER (v5.1).^33^ The HOMER script findMotifsGenome.pl was run using the reference genome FASTA corresponding to the nf-core ATAC-seq genome build. Motifs were identified using the original peak sizes (-size given), and repetitive elements were masked (-mask). Following motif discovery, significantly enriched motifs were annotated across all input regions using annotatePeaks.pl. Motif position weight matrices generated by HOMER excluded redundant “similar” motifs. Motif occurrences were reported in tabular format and as embedded BED coordinates using the -mbed option.

## Data and code availability

RNA-seq and ATAC-seq data generated in this study have been deposited at the Gene Expression Omnibus (GEO) and will be made available upon publication. All data analysis were performed within a containerized environment using Podman. Instructions for setting up the environment and notebooks to run the analysis and reproduce the figures in this article will be shared upon publication via our Github organization (https://github.com/cancerbits) and Zenodo.

Publicly available and annotated single cell RNA-seq data were downloaded as rds objects. The Tabula Muris dataset was downloaded from cellxgene (https://datasets.cellxgene.cziscience.com/2a37a272-6f79-436b-ae11-c1b0b1f293dd.rds),^24^ the immune cell dataset from the Human Cell Atlas was downloaded from Zenodo (https://zenodo.org/records/10197112)^26^ and the iPSC-derived hematopoiesis dataset is available in GEO under the accession number GSE277485.^29^ RNA-seq of sorted tissue-resident macrophages from different tissues established by Lavin et al.^59^ can be found in GEO under the accession number GSE63340. Bulk RNA-seq of sorted tissue-resident macrophages and their precursors established by Mass et al.^3^ was downloaded from the medical epigenomic platform (https://medical-epigenomics.org/papers/mass2016/data/macrophages_bulk_csv.zip).

## Acknowledgements

We would like to thank the Biomedical Sequencing Facility at CeMM for assistance with next generation sequencing, Christoph Frield, from Core Facility Imaging of the Medical University of Vienna, Sophia Lindorfer, Rohit Jain, Shweta Tikoo, and all members of the Farlik and Halbritter labs for their help and advice.

This research was funded in whole or in part by the Austrian Science Fund (FWF) [10.55776/F61]. For open access purposes, the author has applied a CC BY public copyright license to any author accepted manuscript version arising from this submission. Furthermore, the authors would like to acknowledge the following funding sources for their support: Austrian Academy of Sciences (26527 to D.P.), FWF (10.55776/ESP652 to M.P.; 10.55776/PAT1300223 and 10.55776/TAI454 to F.H.), the Federal Ministry of Women, Science and Research and the Ludwig Boltzmann Gesellschaft (LBG) as part of the Clinical Research Groups programme (LBG_KFG_2024_105 to M.F.), and the Alex’s Lemonade Stand Foundation for Childhood Cancer (grant agreement 20-17258; to F.H. and M.F.).

## Author contributions

T.D., F.H. and M.F. planned the study. D.P. and L.E.S performed the experiments. M.P. and F.H. analyzed the data with contributions from D.P., F.A. and MF.. M.F. and F.H. supervised the research. D.P., M.P, F.H. and M.F. wrote the manuscript with contributions from all authors.

## Declaration of generative AI and AI-assisted technologies in the writing process

During the preparation of this work, the authors used ChatGPT-5.5 Plus and Microsoft Copilot (GPT-5) in order to assist with language editing and improve text clarity. After using these tools, the authors reviewed, edited and verified the content as needed and take full responsibility for the content of the manuscript.

---

**Fig. S1.**
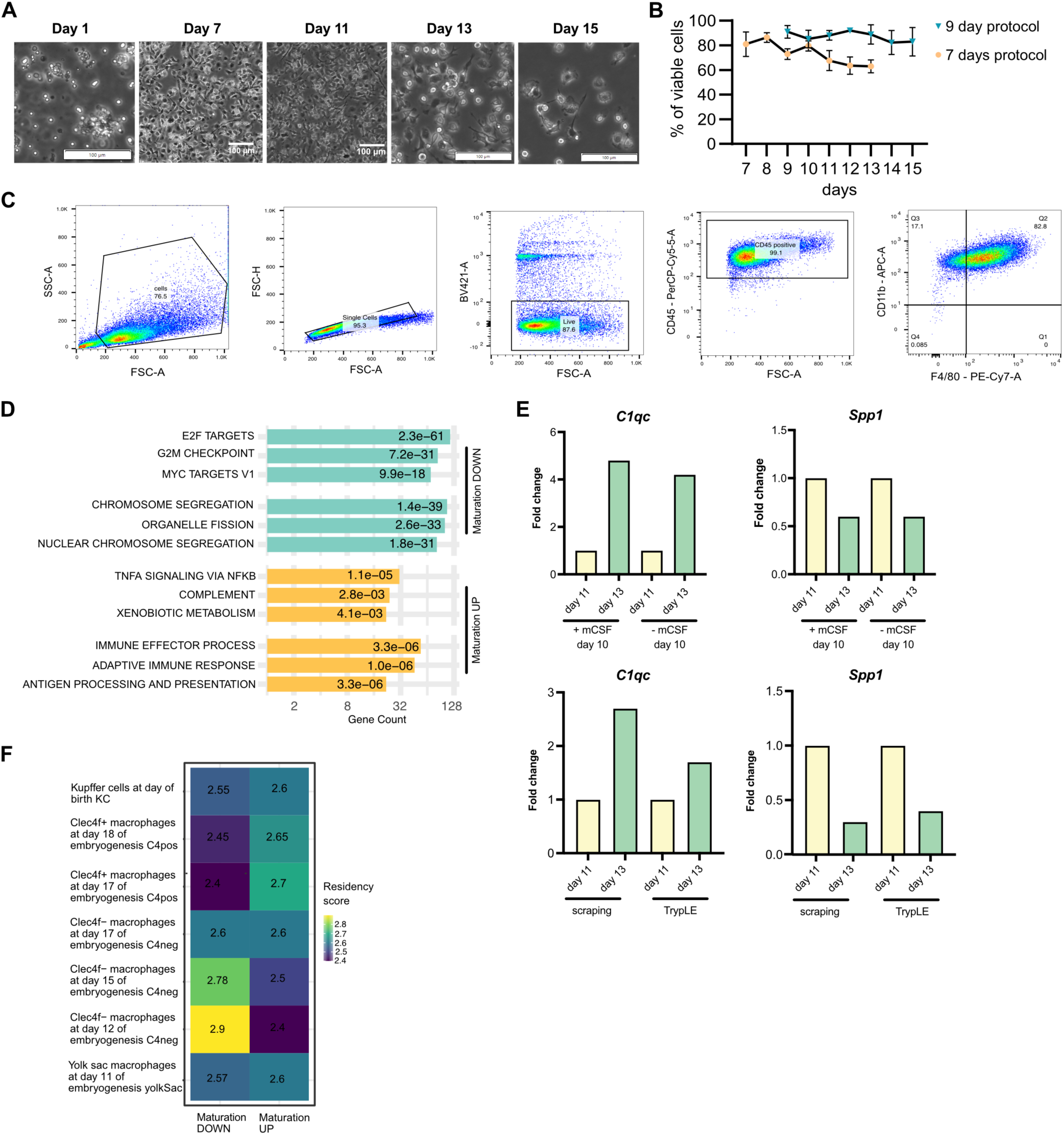
Long-term culture of murine BMDMs recapitulates a core macrophage maturation program with residency-associated features irrespective of origin. A) Representative brightfield images of cells during bone marrow–derived macrophage (BMDM) differentiation after seeding from bone marrow aspirates, captured at days 1, 7, 11, 13, and 15. Scale bar = 100 µm. B) Percentage of viable macrophages (y-axis) over time (x-axis, days) during BMDM differentiation under protocols with M-CSF replenishment until day 7 (orange) or day 9 (blue). Viability was assessed by flow cytometry. C) Flow cytometry gating strategy used to assess BMDM differentiation quality. From left to right: FSC-A/SSC-A gating to select cells; singlet selection using FSC-A/FSC-H; live cell selection based on exclusion of live/dead staining (BV421-A); CD45⁺ cell selection (PerCP-Cy5.5-A); and macrophage identification based on F4/80 (x-axis, PE-Cy7-A) and CD11b (y-axis, APC-A) expression D) Functional enrichment analysis of genes upregulated (yellow) and downregulated (blue) during maturation (day 13 vs day 11), using the Hallmark and Gene Ontology (GO) biological process (C5:BP) gene sets from the Molecular Signatures Database (MSigDB; https://igordot.github.io/msigdbr/). Enrichment was assessed using a one-sided hypergeometric test with false discovery rate correction (Benjamini–Hochberg).^97^ The x-axis indicates the number of enriched genes, and adjusted p values are shown in the labels. The background gene universe corresponds to all genes detected in the RNA-seq dataset. E) qPCR analysis of C1qc (left) and Spp1 (right) expression under different culture conditions. Top: comparison of cultures with or without M-CSF at day 10. Bottom: comparison of cell detachment methods (scraping vs TrypLE enzymatic dissociation). RNA was harvested at day 11 (yellow) and day 13 (green). Expression levels are shown as fold change relative to day 11 within each condition. n = 1 biological replicate. F) Re-analysis of the Bonnardel et al.^25^ bulk RNA-seq dataset of embryonic macrophages and Kupffer cells from day 11 of embryogenesis to day of birth. *ssGSEA* maturation scores were calculated using the *Maturation-UP* and *Maturation-DOWN* signatures established in 1C).

**Fig. S2.**
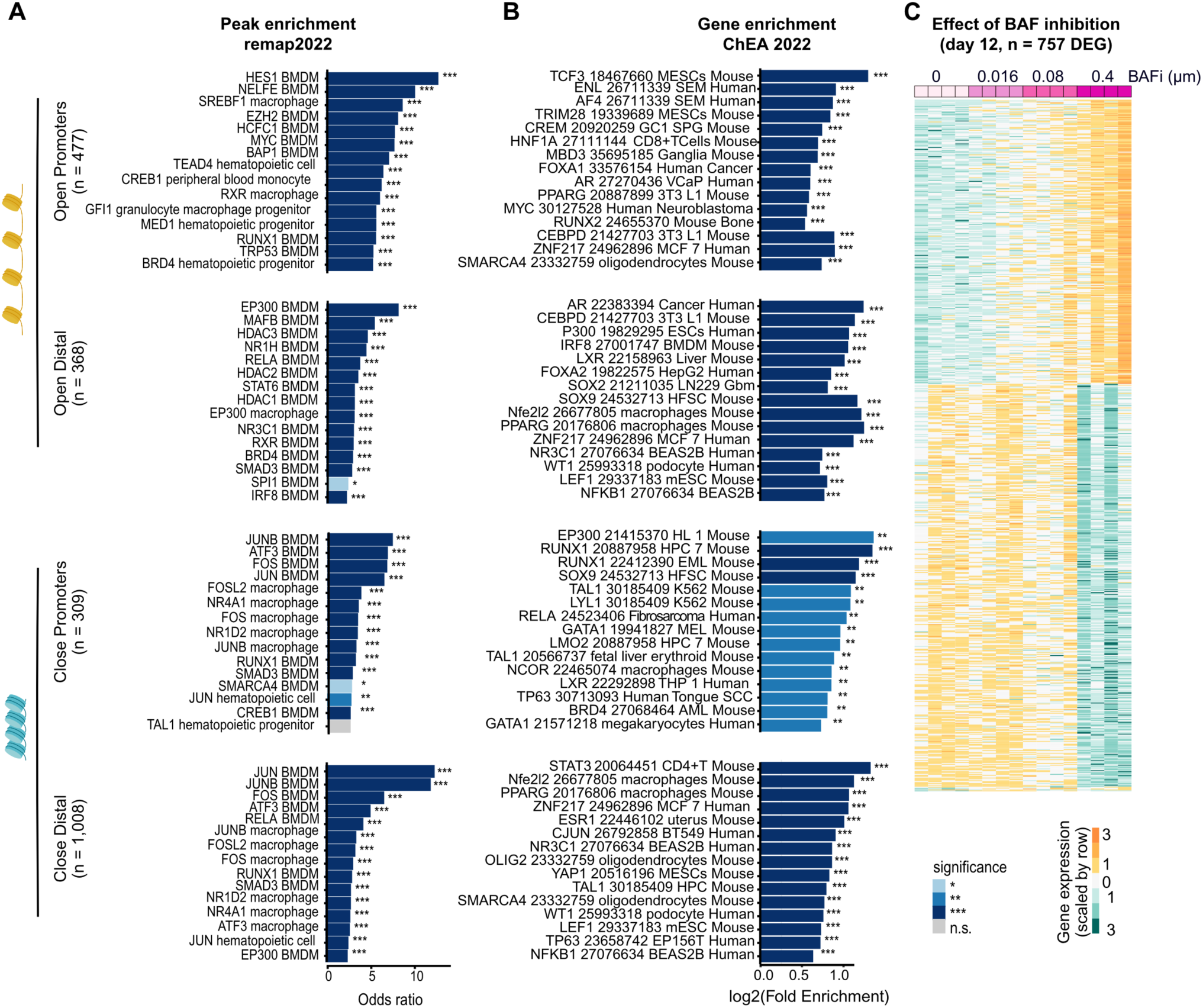
Epigenomic changes during longterm bone marrow–derived macrophage (BMDM) culture. A) Genomic Locus Overlap Enrichment Analysis (LOLA) of differentially accessible regions (DARs) between day 13 and day 11, stratified according to the four clusters defined in A) (open promoter, n = 477 DARs; open distal, n = 368 DARs; closed promoter, n = 309 DARs; closed distal, n = 1,008 DARs). Enrichment was performed against the database of transcriptional regulators peaks from ReMap2022.^36^ Statistical significance was assessed using Fisher’s exact test implemented in LOLA (FDR-adjusted p < 0.05). The x-axis represents the odds ratio, and adjusted FDR values are indicated in the labels (ns, grey; FDR < 0.05, light blue; FDR < 0.005, blue; FDR < 0,0005 dark blue). The background universe corresponds to all accessible regions detected in the ATAC-Seq dataset. B) Functional enrichment analysis of the nearest associated genes to the defined DARs using ChEA gene sets.^96,105^ Enrichment was assessed using a one-sided hypergeometric test with false discovery rate correction (Benjamini–Hochberg).^97^ The x-axis shows the number of enriched genes, and adjusted *P* values are indicated in the labels. The background gene universe corresponds to all genes associated with peak detected in the ATAC-seq dataset. C) Heatmap of differentially expressed genes (DEGs) induced by BAF inhibition, identified by comparing BAF inhibitor–treated samples (BRM/BRG1 ATP Inhibitor-1 - BRM014; 0.016, 0.08, and 0.4 µM; pink gradient from light to dark) to untreated controls. The inhibitor was applied at day 12 and RNA was collected at day 13 (DESeq2 with apeglm LFC shrinkage; adjusted p < 0.05; |fold change| > 1.5). Expression values are scaled by row (turquoise to yellow).

**Fig. S3.**
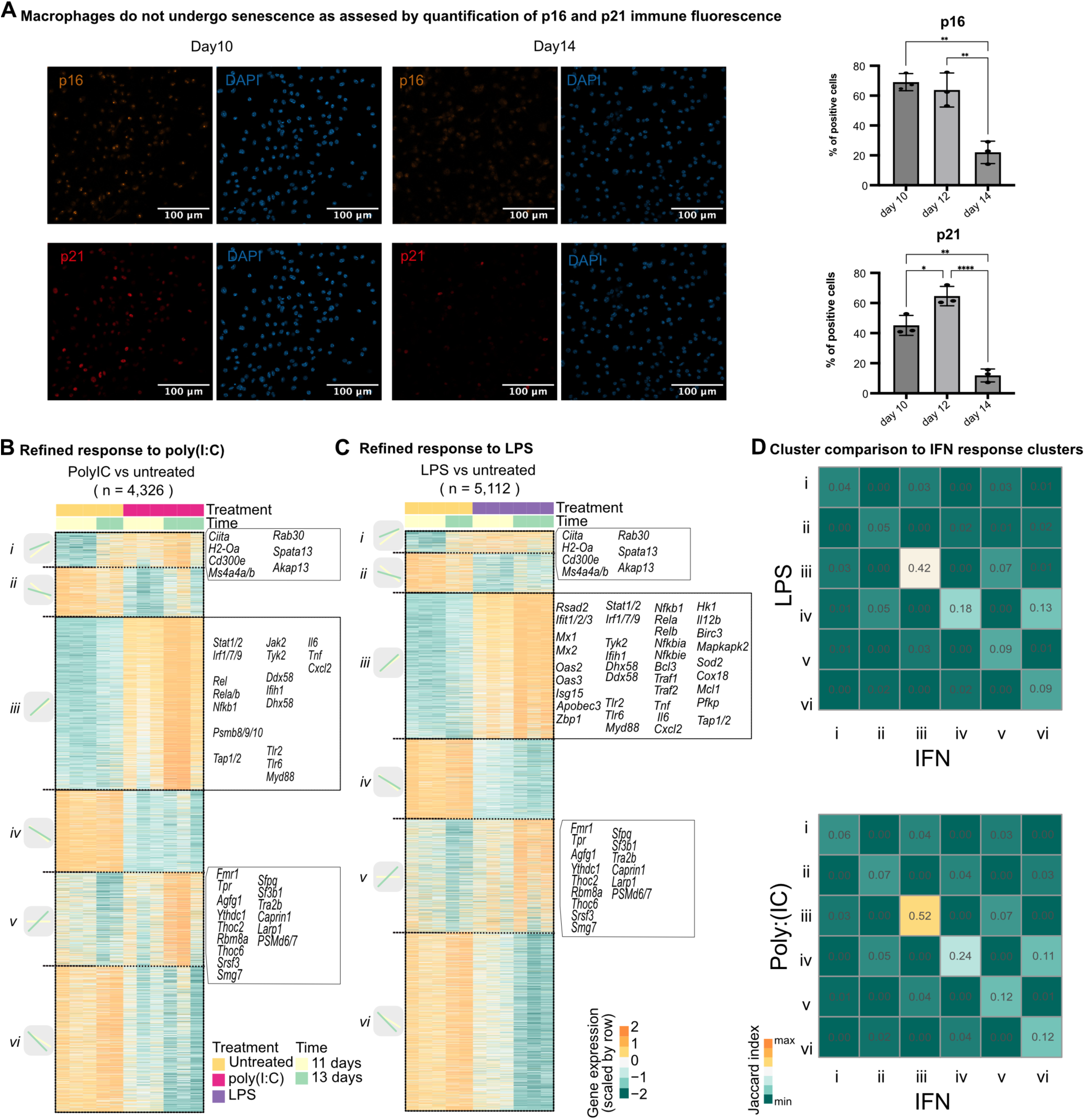
Senescence exclusion and stimulus response profiling after treatment with poly(I:C) and LPS. A) Immune fluorescence staining of senescence markers p16 and p21 in macrophage cultures at day 10 and 14 (left). Scale bar indicates 100µM. Images are representative of 3 biological replicates. Quantification of the immune fluorescence signal measured in macrophages at day 10, 12 and 14. Segmented nuclei from the DAPI channel were quantified using the Cellpose cyto3 model, and object classification was performed in ilastik using the nuclear masks and corresponding raw P16/P21 image data (right). B)-C) Heatmap of differentially expressed genes (DEGs) in response to 4-hour poly(I:C) (B) or LPS (C) treatment in immature (day 11) and mature (day 13) BMDMs (DESeq2 with apeglm LFC shrinkage; adjusted P < 0.05, |fold change| > 1.5). Expression values are scaled by row (from min to max, turquoise to yellow). DEGs are ordered by fold change across conditions. Clusters i-ii represent responsive genes specific to immature BMDMs, respectively up and down regulated; clusters iii-iv represent shared responsive genes, respectively up and down regulated; clusters v-vi contain response genes specific to mature BMDMs. D) Heatmap of pairwise Jaccard indices between LPS and IFN (upper panel) and poly(I:C) and IFN-I (lower panel) response gene clusters (i–vi, as defined in B-C and Fig. 5C). The Jaccard index was calculated as J(A,B)=∣A∩B∣/∣A∪B∣, where A and B represent gene sets from individual clusters. Continuous scale from minimum to maximum overlap (turquoise to yellow), with higher values indicating greater overlap between gene sets.

## List of supplementary tables

Table S1. RNA-seq and ATAC-seq data overview

Table S2. Maturation-related DEGs

Table S3. Maturation-related DARs

Table S4. Results of genomic region overlap analyses (LOLA)

Table S5. Results of motif enrichment analyses (HOMER)

Table S6. Results of geneset overrepresentation analyses (hypeR)

Table S7. BAFi-treatment-related DEGs

Table S8. DEGs related to IFNb, PolyIC, LPS, and memory DEGs

Table S9. LPS vs PolyIC DEGs

Table S10. Results of geneset enrichment analysis (GSEA)

